# Predictive and Experimental Motif Interaction Analysis Identifies Functions of the WNK-OSR1/SPAK Pathway

**DOI:** 10.1101/2024.06.26.600905

**Authors:** Clinton A Taylor, Ji-Ung Jung, Sachith Gallolu Kankanamalage, Justin Li, Magdalena Grzemska, Ankita B. Jaykumar, Svetlana Earnest, Steve Stippec, Purbita Saha, Eustolia Sauceda, Melanie H. Cobb

## Abstract

The WNK-OSR1/SPAK protein kinase signaling pathway regulates ion homeostasis and cell volume, but its other functions are poorly understood. To uncover undefined signaling functions of the pathway we analyzed the binding specificity of the conserved C-terminal (CCT) domains of OSR1 and SPAK to find all possible interaction motifs in human proteins. These kinases bind the core consensus sequences R-F-x-V/I and R-x-F-x-V/I. Motifs were ranked based on sequence, conservation, cellular localization, and solvent accessibility. Out of nearly 3,700 motifs identified, 90% of previously published motifs were within the top 2% of those predicted. Selected candidates (TSC22D1, CAVIN1, ATG9A, NOS3, ARHGEF5) were tested. Upstream kinases WNKs 1-4 and their close relatives, the pseudokinases NRBP1/2, contain CCT-like domains as well. We identified additional distinct motif variants lacking the conserved arginine previously thought to be required, and found that the NRBP1 CCT-like domain binds TSC22D1 via the same motif as OSR1 and SPAK. Our results further highlight the rich and diverse functionality of CCT and CCT-like domains in connecting WNK signaling to cellular processes.

## INTRODUCTION

The WNK [with no lysine (K)] signaling cascade is comprised of the four WNK kinases (WNKs 1-4) and their downstream substrate kinases, oxidative stress responsive 1 (OSR1) and SPS/STE20-related proline-alanine-rich kinase (SPAK). Ancient and pleiotropic kinases, WNKs are so named due to their unique placement of a catalytic lysine required for ATP binding compared to other members of the eukaryotic kinase family (Verissimo and Jordan, 2001; Xu et al., 2000). Mutations that cause overexpression of WNK1 and WNK4 cause an inherited form of hypertension, pseudohypoaldosteronism type II, thereby linking the WNK pathway to control of blood pressure (Wilson et al., 2001). Further demonstrating its importance in the cardiovascular system, disruption of the WNK1 gene in mice revealed its requirement in the endothelium for cardiovascular development (Xie et al., 2009).

Subsequent studies found that WNKs interact with and activate OSR1 and SPAK by phosphorylation, enabling OSR1 and SPAK to phosphorylate a variety of substrates, some of which are associated with ion transport and cell volume regulation (Anselmo et al., 2006; Darman and Forbush, 2002; de Los Heros et al., 2014; Dowd and Forbush, 2003; Gagnon et al., 2007a; Moriguchi et al., 2005; Pacheco-Alvarez et al., 2012; Piechotta et al., 2003; Piechotta et al., 2002; Ponce-Coria et al., 2008; Richardson et al., 2008; Richardson et al., 2011; Taylor et al., 2018; Vitari et al., 2005; Vitari et al., 2006). Although the actions of the WNK-OSR1/SPAK pathway as a central regulator of ion homeostasis and cell volume has been extensively studied, its other functions are still poorly understood. Mutations in WNK1 can also cause a form of hereditary and sensory neuropathy (HSN2) (Shekarabi et al., 2008). WNKs are ancient kinases that have adapted to regulate many cellular processes. Previous studies have suggested inputs of WNK1 to metabolic pathways and others have identified single nucleotide polymorphisms in WNK genes not only in hypertensive disease but also, for example, in neurological syndromes including autism and schizophrenia (Arion and Lewis, 2011; Henriques et al., 2020; Mendes et al., 2010; Prisco et al., 2021; Qiao et al., 2008; Ramoz et al., 2008; Shekarabi et al., 2017; Siew and O’Shaughnessy, 2013; Tobin et al., 2005; Vavolizza et al., 2012; Wang et al., 2009). WNK kinases have been linked to numerous cellular activities suggesting they are multifaceted in the processes they regulate. Examples include but are not limited to autophagy, cell migration, insulin signaling, fat metabolism, neuronal signaling, and immune system function (Gallolu Kankanamalage et al., 2016; Hayward et al., 2023; Jaykumar et al., 2021; Köchl et al., 2016; Mendes et al., 2010; Shekarabi et al., 2008; Sun et al., 2021).

Our goal was to deduce understudied signaling functions of the pathway by analyzing the capacity of the kinases OSR1 and SPAK to recognize short protein sequence motifs in the proteome through their conserved C-terminal (CCT) domains. Because SPAK and OSR1 are proteins of average size with well-defined domains compared to WNKs, which are large (WNK1, most common 2126 and 2384 residues) with much low complexity sequence, we surmised that we could gain valuable insights into WNK signaling by investigating proteins that bind SPAK and OSR1.

We first characterized the binding specificity of the CCT domains of OSR1 and SPAK; these domains interact with the core consensus protein sequences R-F-x-V/I which are found in WNKs, substrate ion cotransporters, and other proteins (Delpire and Gagnon, 2007; Gagnon et al., 2007a; Moriguchi et al., 2005; Piechotta et al., 2003; Piechotta et al., 2002; Taylor and Cobb, 2022; Villa et al., 2007). Motifs with an alternative placement of the arginine (R-x-F-x-V/I) also bind to the CCTs, as we previously showed in a subset of potassium channels (Taylor et al., 2018). Here, after noting that identities of the flanking residues surrounding the core motifs contribute significantly to overall binding, we used the specificity data to identify all possible interaction motifs in human proteins, and ranked them by predicted interaction strength. After removing likely false-positives, motifs were scored based on conservation, localization, motif sequence, and solvent accessibility. Of the 27 previously validated motifs, 24 ranked in the top 2% of all identified motifs, implying the ranking methodology worked well. We evaluated a subset of candidate interactors in vitro and including some in cellular contexts. Additionally, WNKs themselves each contain two CCT-like domains and the WNK related pseudokinases NRBP1/2 each contain one CCT-like domain. We show that the NRBP1 CCT-like domain can interact with the same motif as SPAK/OSR1.

Our results have deciphered additional features of CCT-motif interactions that expand knowledge of the already large potential of the CCT and CCT-like domains to connect WNK signaling to cellular regulatory events. OSR1 and SPAK protein interactions themselves account for a rich and understudied source of regulatory power for the WNK pathway.

## RESULTS

### Core motif and flanking residues participate in SPAK/OSR1 CCT interaction specificity

The CCT domains of OSR1 and SPAK are small (∼90 residues) modular domains found at the C-terminus of both kinases comprised of a β-sheet and two α-helices (**Figure 1A,B)**. Structural data are also available for some of the related domains of which two are found in each WNK (CCTL1, CCTL2), for example. The crystal structures of the OSR1 CCT bound to a GRFQVT hexapeptide (PDB ID: 2V3S) and the related domain, WNK2 CCTL1, bound to a longer WNK1-derived RRFxV-containing peptide that engages the residues R-x-F-x-V (PDB ID: 6FBK, Structural Genomics Consortium) both indicate that residues around the core motif binding pocket associate in part through β-strand addition, whereby backbone atoms from a β-strand in one binding partner interact with backbone atoms in a β-strand from the other partner in order to impart additional binding affinity **(Figure 1B)** (Pinkas et al., 2017; Remaut and Waksman, 2006; Villa et al., 2007). The result of this mode of binding is that adjacent residues within and around the core motif alternate direction.

**Figure 1.**
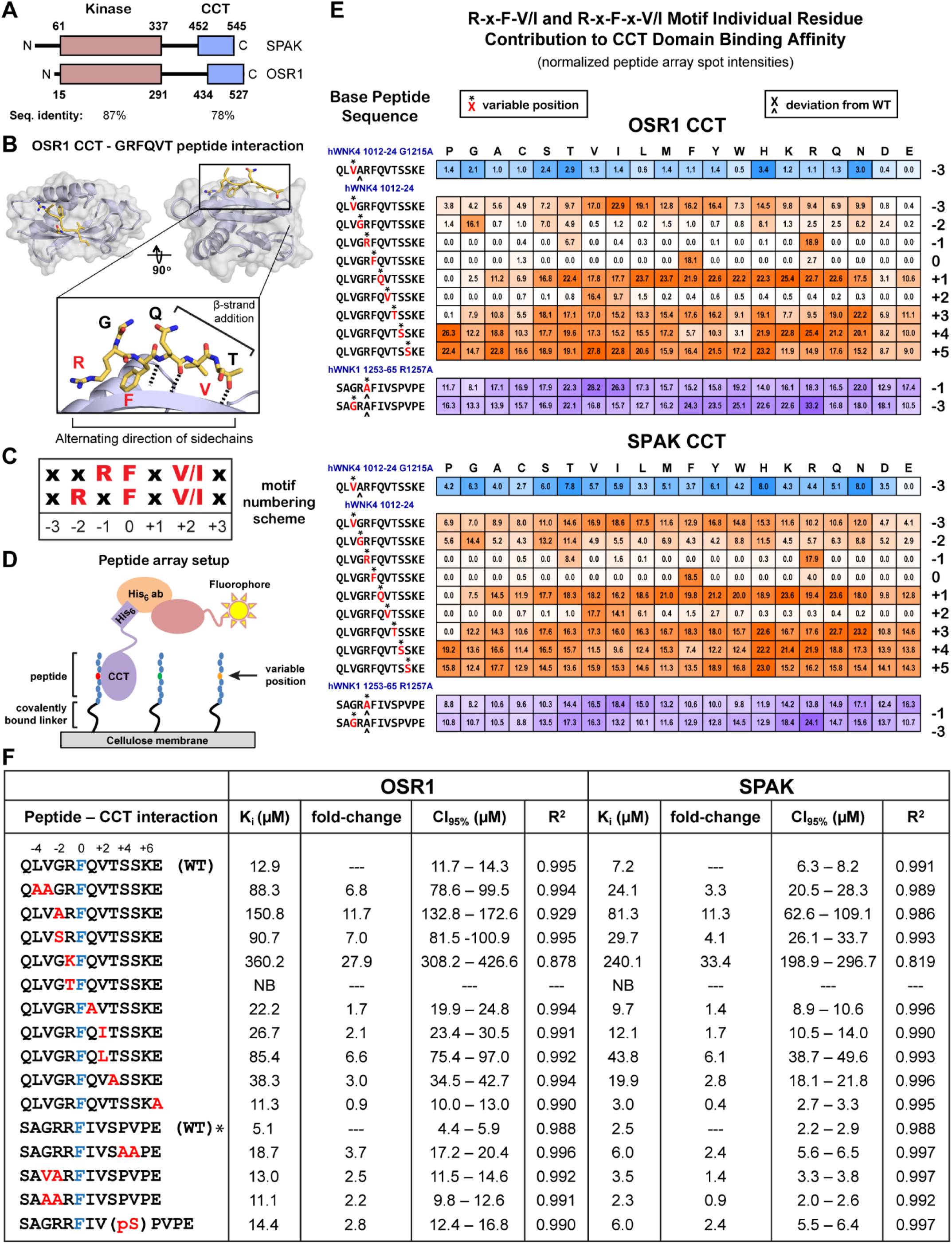
Contributions to OSR1 and SPAK CCT domain binding by residues within and around core RFxV/I and RxFxV/I motifs. **(A)** Schematic representation of the paralogous kinases SPAK (STK39) and OSR1 (OXSR1). Both have highly conserved N-terminal kinase domains and C-terminal CCT domains. CCT domains interact with R-F-x-V/I and R-x-F-x-V/I motifs. **(B)** Crystal structure of the complex between the OSR1 CCT domain and a WNK1-derived hexapeptide, GRFQVT (PDB ID: 2V3S). The peptide interacts partially by acting as a terminal β-strand (β-strand addition), and as a result, the sidechains alternate direction suggesting adjacent residue identities do no strongly influence binding of neighboring residues. **(C)** Numbering scheme used in this paper to describe positions of residues along motif. Phenylalanine, F, is fixed at the zero position. **(D)** Cartoon representation of peptide array setup. Peptides are covalently linked to a membrane. Each spot represents a unique peptide where a single position within the peptide is varied to every amino acid. Purified His_6_-CCT domains bind to the peptides and fluorescent antibodies binding the His_6_ tag provide quantification of binding strength. **(E)** Quantification of peptide array spot intensities. Original peptide array images shown in Supp. Fig. 1. Asterisk (*) above the red residue indicates the position varied to every amino acid. Caret (^) below the residue indicates a site in the base peptide that deviates from the wild type sequence. Number to right indicates motif position mutated. Darker color indicates stronger signal. In the top (blue) row the glycine in the peptide hWNK4 (human WNK4) 1012-1024 is mutated to alanine (G1215A) in order to analyze the binding of the adjacent amino acid in the presence of a less flexible residue. The middle (orange) group represents the wild type hWNK4 sequence. The bottom (purple) group analyzes the binding contributions of the residues surrounding the arginine in the alternative motif R-x-F-x-V/I in the peptide sequence hWNK1 1253-1265 R1257A. The wild type hWNK1 sequence contains both motifs (R-R-F-x-V/I). n=2 peptide arrays per CCT domain normalized by the mean intensity of the array and reported as the average. Ranges are reported in **Supp. Fig. 2. (F)** Affinity determined by fluorescence anisotropy peptide competition assays. Unlabeled peptides at various concentrations displace labeled peptide (NH_3_^+^-NLVGRF-[DAP-FAM]-VSPVPE-COO^−^] [diaminopropionic acid (DAP)). Labeled peptide held constant at 25 nM, SPAK CCT and OSR1 CCT are constant at 1.5 μM and 3.0 μM, respectively. Red letters are mutated relative to the wild type (WT) sequence. Blue letter is motif phenylalanine (position 0). Top group of peptides is hWNK4 1012-1024 and bottom is hWNK1 1253-1265. Affinity reported as inhibition dissociation constant (K_i_, µM). Upper and lower limits of 95% confidence interval indicated. Fold-change decrease in affinity (fold-change increase in K_i_) relative to wild type indicated. Goodness of fit reported as R-squared (R^2^). All measurements are n=3. Fit curves and fluorescent probe affinity measurement can be found in **Supp. Fig. 3.** GraphPad Prism 10 with one site fit to K_i_ model used for data analysis. Asterisk (*) indicates results previously published (Taylor et al., 2018).

Therefore, we decided to systematically examine the effects of mutating each residue to every other amino acid around the core motif because presumably the interactions with the CCT domain of the adjacent residues to the residue being mutated should be minimally impacted due to the alternating directions **(Figure 1B)**. To describe these motif-containing peptides we designated F in the motif as the “0” position, residues N-terminal as negative (-1, -2, etc…) and residues C-terminal as positive (+1, +2, etc…). **(Figure 1C)**. Phenylalanine at the 0 position is likely the most invariant and structurally fixed residue. To accomplish the systematic analysis we used peptide arrays in which the sequence of a base peptide is mutated at each residue position to all other amino acids **(Figure 1D,E; Supp. Figures 1,2)**. Bacterially-expressed His_6_-tagged SPAK and OSR1 CCT domains were incubated with a cellulose membrane containing covalently-attached peptides that were synthesized directly on the membrane. Residual protein was washed off, the membrane was probed with a primary anti-His_6_ antibody, a secondary fluorescent antibody, and imaged using a fluorescence imaging system to quantify the amount of CCT domain remaining bound to each peptide spot **(Figure 1D)**.

We chose three base peptides for analysis **(Figure 1E)**. The wild-type WNK4 sequence QLVGRFQVTSSKE (1012-1024) was used as the primary sequence for analysis, WNK4 QLVARFQVTSSKE (1012-1024 G1215A) was used to interrogate the role of the hydrophobic valine at the -3 position in the presence of the less flexible alanine at the -2 position, and WNK1 SAGRAFIVSPVPE (1253-65 R1257A, wild-type sequence: SAGRRFIVSPVPE) was used to investigate the impact on binding of residues around the alternatively placed arginine in the -2 position in R-x-F-x-V/I motifs by mutating the arginine at the -1 position to either every other amino acid or to alanine when mutating residues at the -3 position **(Figure 1C-E; Supp. Figure 1,2)**. Overall, the binding specificities of the SPAK and OSR1 CCT domains are remarkably similar as might be expected considering the strong conservation of residues within the primary motif binding pocket (Elvers et al., 2022; Taylor and Cobb, 2022; Villa et al., 2007).

For the WNK4 peptide, glycine is the most preferred residue at the -2 position but there is also a preference for S, T, H, and N in OSR1 and SPAK, and additionally, M and W for only SPAK **(Figure 1E)**. Variable binding due to changes in this residue are more pronounced in the OSR1 CCT compared to SPAK, perhaps due to the higher overall affinity of SPAK for these peptides **(Figure 1F)**. In addition, the presence of either glycine or alanine at the -2 position drastically alters the preference of residues at the -3 position for both OSR1 and SPAK, as might be expected based on the flexibility of glycine. Both SPAK and OSR1 have overwhelmingly strong preferences for the core motif residues, as is to be expected; the preference for threonine at the -1 position is an artifact as determined by fluorescence anisotropy described later **(Figure 1F)**. Additionally, proline is strongly disfavored in and around the core motif, as is to be expected since it would disrupt β-strand addition. The crystal structure of OSR1 bound to a GRFQVT peptide indicates the glutamine at the +1 position points out into solvent, and likewise we observe that there is not a strong preference at this position although small or negatively charged sidechains are generally more disfavored. At the +3 to +5 position some residues are clearly favored over others, but nothing is as striking as alterations in the core motif residues. For example, at the +4 position there is clear preference for basic residues or proline, while bulkier hydrophobic side chains are disfavored. Interestingly, of the 27 known, experimentally validated motifs, proline is present at the +4 position in six while a basic residue is present in five indicating an agreement between our peptide array results and previously identified motifs **(Figure 2B)**.

**Figure 2.**
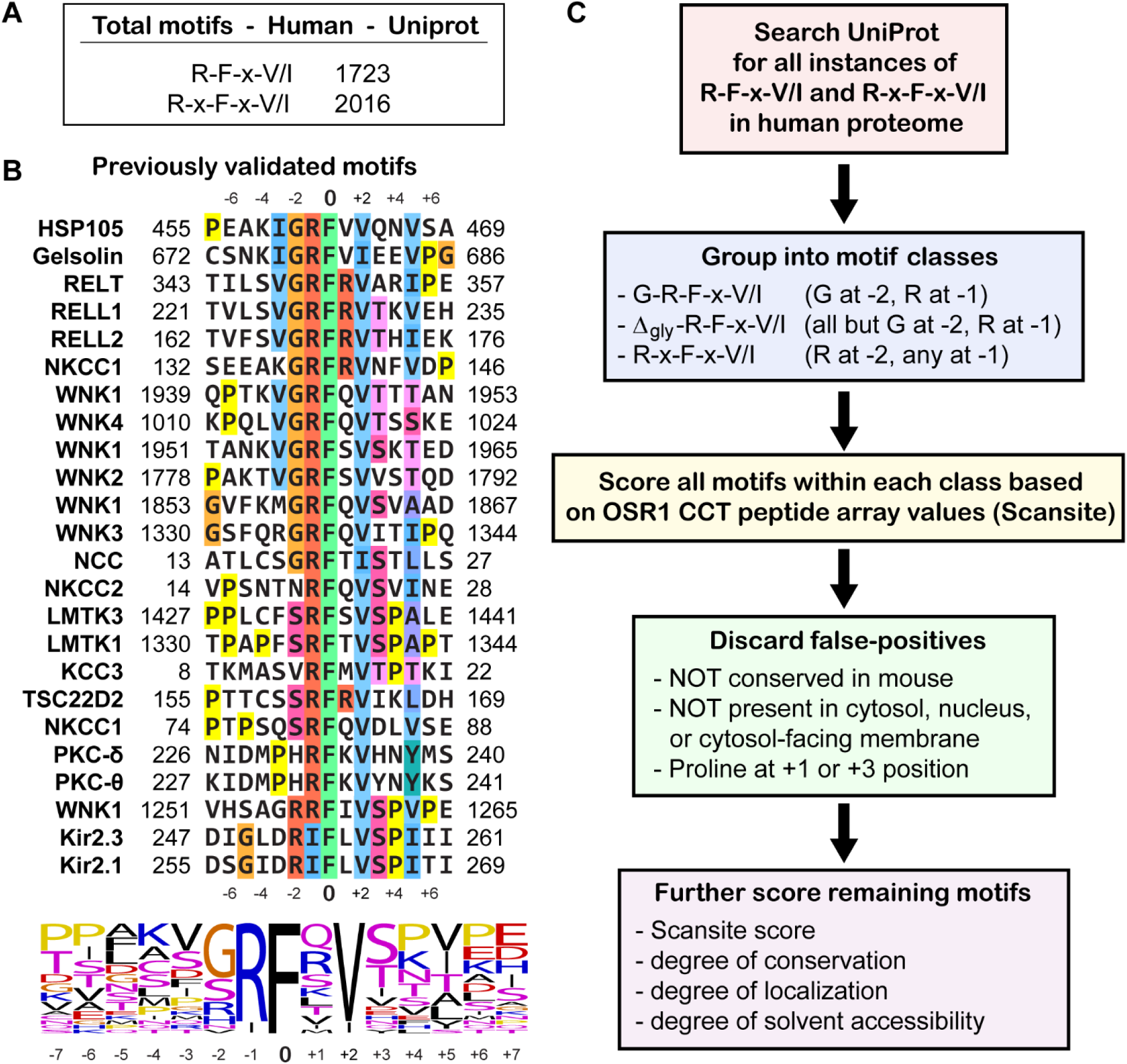
Summary of protein interaction prediction process. **(A)** Total motifs found in initial search depending on core motif type (R-F-x-V/I or R-x-F-x-V/I). **(B)** Multiple sequence alignment of all experimentally identified motifs (Anselmo et al., 2006; Delpire and Gagnon, 2007; Gagnon and Delpire, 2010; Gagnon et al., 2006; Gagnon et al., 2007a; Gagnon et al., 2007b; Jaykumar et al., 2024; Li et al., 2016; Li et al., 2004; Moriguchi et al., 2005; Piechotta et al., 2003; Piechotta et al., 2002; Polek et al., 2006a; Polek et al., 2006b; Richardson et al., 2008; Smith et al., 2008; Taylor et al., 2018; Villa et al., 2007; Vitari et al., 2006). Results generated with Clustal Omega (Sievers et al., 2011). Sequence logo generated with WebLogo (Crooks et al., 2004) **(C)** Flowchart describing entire process used in this paper to predict CCT domain interactions with motifs. Human motifs comprising three different classes were identified in the UniProt database and scored using the peptide array data with Scansite3 (Obenauer et al., 2003). Then false-positives were removed. Finally, motifs were further scored based on conservation, localization, solvent accessibility, and the initial Scansite score. A detailed flow-chart with scoring information can be found in **Supp. Fig. 4.**

The identity of residues around the -2 arginine in the WNK1 (R-x-F-x-V) mutant peptide have a less pronounced effect **(Figure 1E)**. At the -1 position there is a slight preference for branched chain amino acids, threonine, asparagine, and glutamic acid. At the -3 position the strongest preference is for arginine suggesting that arginine further out from the core motif may be able to replace the arginine at the +1 position much like the arginine in R-x-F-x-V/I motifs does, or at least provide additional binding energy. Yet, the entropic costs to binding likely increase with an arginine placed further away from the centrally fixed phenylalanine. In addition, the preferences for SPAK and OSR1 at the -3 position diverge slightly aside from the strongest preference for arginine.

Because peptide arrays only provide a semi-quantitative measure of relative binding affinity, we quantified binding to a select group of sequences utilizing the same two base peptides derived from WNK4 (QLVGRFQVTSSKE) and WNK1 (SAGRRFIVSPVPE). Fluorophore labeled peptides are more difficult to produce and can lead to artifacts due to the presence of a conjugated fluorophore so we chose to utilize peptide competition to measure binding affinity, where unlabeled peptide is used to compete with labeled peptide. This was accomplished via plate-based fluorescence anisotropy measurements where SPAK and OSR1 CCT domains and labeled probe peptide were held constant while titrating the unlabeled peptide to compete off the labeled probe in order to measure the K_i_, the inhibitory constant, which is a type of equilibrium dissociation constant (K_d_) **(Figure 1F, Supp. Figure 3A)**. Fluorescence anisotropy (also known as polarization) measures how fast a fluorophore rotates in solution. A bound fluorophore-labeled peptide will rotate slower and have a higher anisotropic signal. To optimize the assay, we tested several fluorescent probe peptides and ultimately chose the probe NH_3_^+^-NLVGRF-[DAP-FAM]-VSPVPE-COO^-^, which combines the N-terminus of the WNK4 peptide and the C-terminus of the WNK1 peptide with an internal diaminopropionic acid conjugated to the fluorophore FAM and the N-terminal glutamine mutated to asparagine for enhanced stability. We reasoned that an internal fluorophore with a short linker would experience higher anisotropy when bound and thus give a better signal-to-noise ratio. A shorter WNK4 peptide bound with weak affinity and an N-terminally labeled hybrid peptide bound with higher affinity but was not stable over time **(Supp. Figure 3B, 3C)**. One limitation of our assay is that peptide affinities much higher than the K_D_ of the probe peptide (SPAK: 0.67 µM, OSR1: 1.5 µM) cannot be reliably measured. However, none of the peptides measured appeared to cross this threshold, which would appear as a sharper than expected drop in anisotropic signal during peptide titration **(Supp. Figure 3A)**.

Overall, we consistently observed SPAK CCT as having several-fold higher affinities for peptides compared to OSR1, which likely explains the stronger overall binding observed by peptide array for SPAK **(Figure 1E; Supp. Figures 1,2)**. However, the measurements are derived from purified CCT domains and short, 13-mer peptides, both of which may affect affinities compared to what might be observed using full-length proteins **(Figure 1F)**.

For the WNK4 wild type peptide, the K_i_ for OSR1 and SPAK CCT domains were 12.9 µM and 7.2 µM, respectively. Mutation of the branched chain amino acids at the -4 and -3 positions to alanine, and glycine to alanine or serine at the -2 position resulted in a significant loss of affinity suggesting the importance of these residues in the interaction. Mutation of the -1 arginine to lysine or threonine effectively prevented the interaction, contrary to what had been observed in the peptide array for threonine, suggesting it was an artifact **(Figure 1E,F)**. Substitution of glutamine with alanine at the +1 position had a minimal effect supporting the notion that this residue pointing into solvent makes its identity of less importance (Villa et al., 2007). Interestingly, valine to isoleucine at the +2 position also had only a modest effect on affinity, while substitution with leucine led to a more pronounced decrease in affinity. Threonine to alanine at the +3 position led to a modest decrease in affinity, which has been predicted based on the crystal structure of the OSR1:GRFQVT peptide complex (Villa et al., 2007). Finally, the mutation of the most C-terminal amino acid in the peptide at the +7 position led to almost no change in OSR1 but a 2.5-fold increase in affinity for the SPAK CCT suggesting residues that interact outside of the primary binding pocket may provide some selectivity for one CCT domain over the other.

For the WNK1 peptide, the K_i_ for OSR1 and SPAK CCT domains were 5.1 µM and 2.5 µM, respectively **(Figure 1F)**, which are similar to results we published previously when first describing the R-x-F-x-V/I motif variant (Taylor et al., 2018), but we have reanalyzed the data utilizing updated software. Mutation of proline and valine to alanines at the +4/+5 positions led to a several fold decrease in affinity. Substitution of -3 glycine and -2 arginine with either valine-alanine or alanine-alanine only resulted in a small loss of affinity implying this glycine is not as important as the one found in the WNK4 motif and that R-F-x-V as a core motif is sufficient for binding, while double arginines at -2/-1 might enhance affinity compared to single arginines. Finally, we also tested the effects of serine phosphorylation at the +3 position since a previous report had indicated that phosphorylation at this position decreased binding affinity by orders of magnitude for a GRFQVpT hexapeptide (Villa et al., 2007). Using the longer WNK1-derived peptide we observed only a modest approximately 2.5-fold decrease in affinity, but the results do imply phosphorylation at the +3 position can modulate binding affinity.

Altogether our peptide array and fluorescence anisotropy results suggest that residues outside the core motif contribute significantly to interaction specificity. In some cases, effects on binding affinity may be due to the presence of one or two strongly favored/disfavored amino acids, in other cases modest, additive effects of multiple residues may be required to provide the necessary specificity, and all these effects may also be context dependent; based on the sequence of adjacent or semi-adjacent residues.

### Application of CCT domain specificity analysis to predict potential interactors

We applied our systemic accounting of OSR1 and SPAK CCT domain specificity determinants to identify and sort all instances of R-F-x-V/I and R-x-F-x-V/I in the human UniProt database by predicted interaction affinities (Uniprot-Consortium, 2015). There are over 3,700 instances of these motifs found in UniProt **(Figure 2A)**. Multiple sequence alignment of all known, previously validated motifs suggest certain partly conserved motif signatures such as -3 valine, -2 glycine/serine, and +3 serine/threonine, but since only 27 motifs have been experimentally verified, not enough sequence space is covered to reliably rank motifs **(Figure 2B)**.

First, we generated scoring matrices utilizing the OSR1 CCT peptide array data **(Figure 1E)**, and using three distinct motif classes. G-R-F-x-V/I and Δ_gly_-R-F-x-V/I (all residues but G at the -2 position) were utilized owing to the large number of motifs found to contain glycine at -2, but also taking into account the dependence on the glycine at -2 must be context dependent since our WNK1 peptide bound with higher affinity than the WNK4 peptide but did not contain a glycine at -2. The third class were the R-x-F-x-V/I motifs, which contain an alternative core motif **(Figure 2C, Supp. Figure 4)**.

We utilized the Scansite3 web server to identify and score all identified motifs based on our input scoring matrices **(Figure 2C, Supp. Figure 4, Supp. Table 1)**. Nearly all of the previously validated motifs clustered near the very top of our lists ranked by predicted motif binding affinity providing an internal metric that indicated our initial method of ranking based solely on sequence worked remarkably well **(Figure 3A,B)**. One limitation of our use of Scansite is that motifs where the central phenylalanine of the motif is within six residues of the N-or C-termini of the protein were not included due to differences in scoring. However, these motifs were still listed for reference **(Supp. Table 1)**.

**Figure 3.**
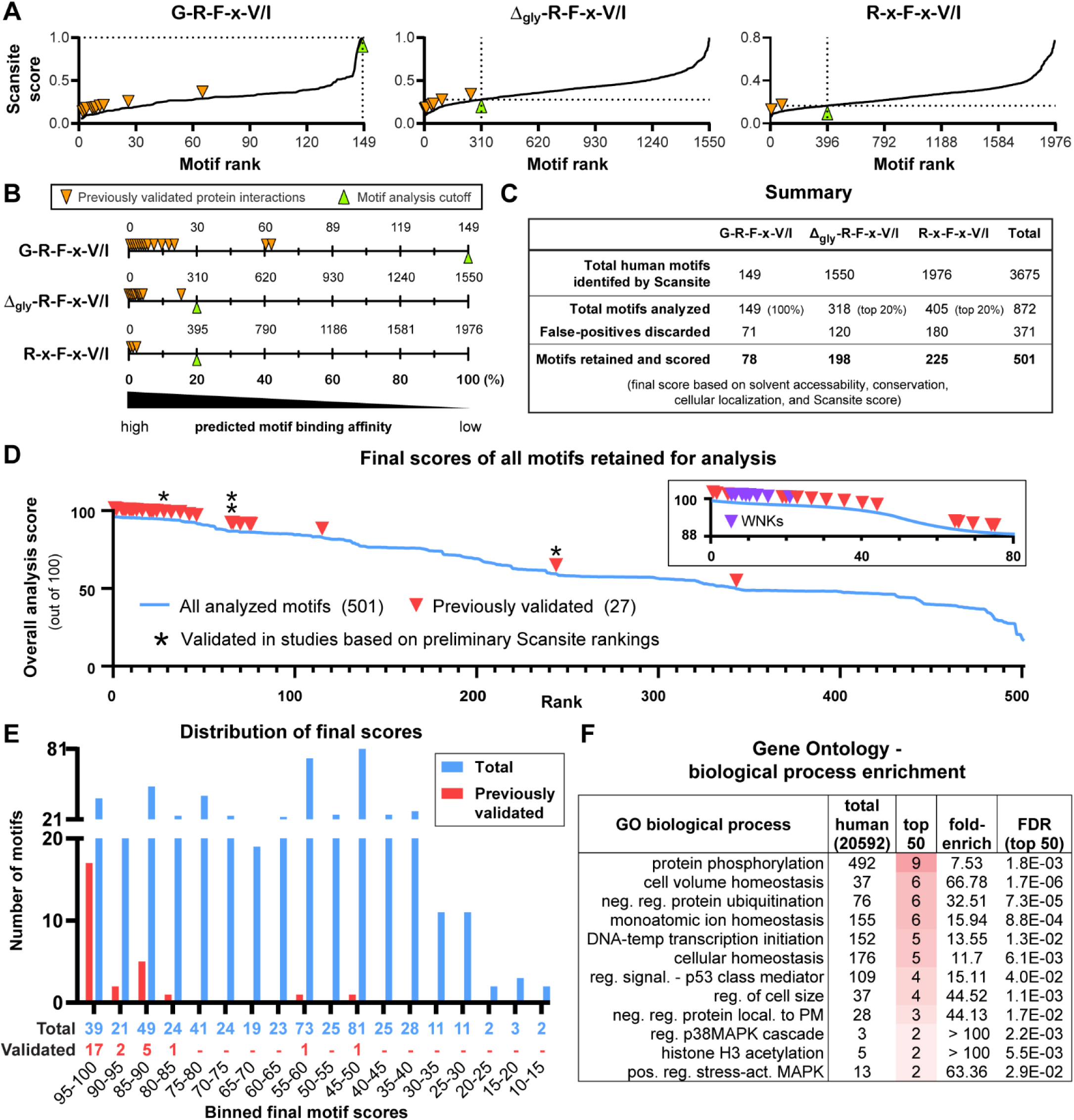
SPAK and OSR1 motif interaction prediction results summary. **(A)** Plots comparing scores of all identified motifs within each motif class found using the Scansite3 web server (Obenauer et al., 2003). A lower score indicates a higher predicted binding affinity. Orange rectangles indicate previously experimentally validated interactions (direct binding or inferred based on other experimental results). Dashed lines indicate cut-offs used for further motif analysis (false-positive filtering and subsequent scoring of remaining motifs). **(B)** Similar to (A) but without scores included on y-axis for clarity. Note that previously validated motifs all cluster at the top of the ranking; indicating that initial motif scoring based on peptide array results was successful. **(C)** Overall comparison of how previously validated motifs were filtered and further scored based on conservation, localization, solvent accessibility, and initial Scansite score. Note that all motifs that had been experimentally validated were retained after. **(D)** Overall scores of all motifs that were retained and subjected to final round of scoring. Blue line indicates all 501 scored motifs. Red triangles indicate positions of all 27 previously validated motifs. Asterisks (*) indicate motifs initially identified in our preliminary Scansite3 analysis that led to publication of validated interactions prior to publication of this study (Jaykumar et al., 2024; Taylor et al., 2018). Inset graph depicts motifs ranked 1 to 80 for clarity. **(E)** Histogram depicting distribution of scores where each bin represents total number of motifs (blue) and previously validated motifs (red) within each bin. Bin size corresponds to all motifs within 5 point intervals. Total numbers of motifs in each bin are listed below. **(F)** Gene ontology (GO) analysis of the top 50 motifs identified in the analysis.

**Table 1.**
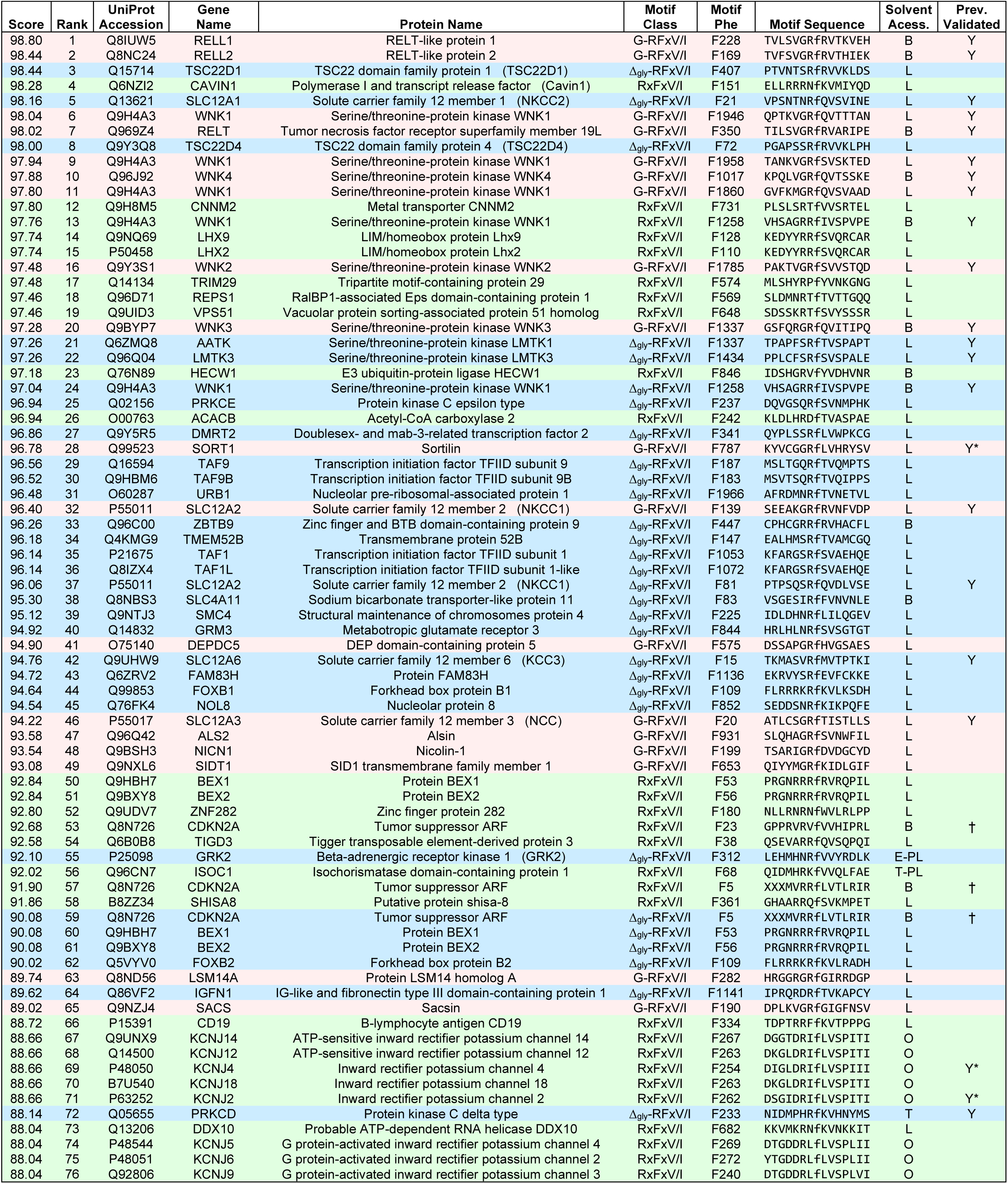
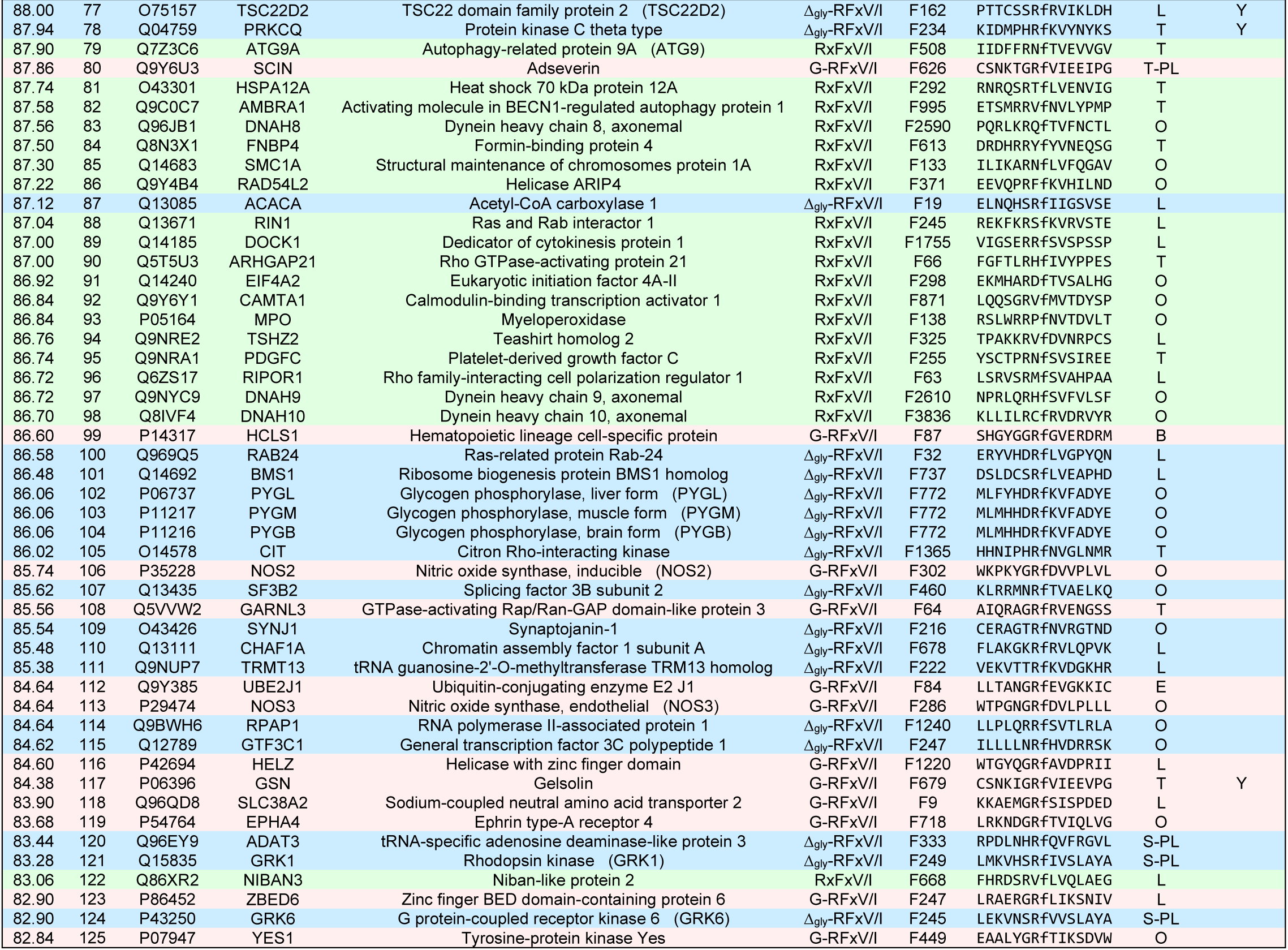
Highest ranked predicted interactions from analyzed motifs. Motifs in table are color-coded by motif class. Score refers to overall score of final analysis and not Scansite score. Score was used to rank where higher scores indicate higher probability of interactions. Position of F refers to “0” position of phenylalanine in the identified motif (lowercase f in sequence) within the protein sequence found in the UniProt database (Uniprot-Consortium, 2015). “Y” in previously validated column indicates previously experimentally validated interaction (direct binding or inferred based on other experimental results). Asterisk (*) next to Y indicates motifs initially identified in our preliminary Scansite3 analysis that led to publication of validated interactions prior to publication of this study (Jaykumar et al., 2024; Taylor et al., 2018). Sequence annotation for protein “Tumor suppressor ARF” (indicated by †) changed during analysis but result did not affect score. “Solvent Access.” refers to motif solvent accessibility as determined by examination of AlphaFold2 models, or when model is not available, experimentally determined structures of protein or structurally-related protein (Jumper et al., 2021; Varadi et al., 2021). Codes for solvent accessibility: L: loop, B: β-hairpin, E: packed β-hairpin, T: terminal β-sheet strand, O: packed loop, S: packed terminal β-sheet strand, and -PL: partial loop. Full details on scoring and entire list of motifs can be found in Methods, **Supp. Figure 4**, and **Supp. Table 1**.

### Identification of strongly predicted interacting proteins following refinement

To further refine our prediction of strong SPAK/OSR1 CCT interaction motifs, we took into account the Scansite score combined with additional factors indicated below **(Figure 2C**, **Figure 3A,B, Supp.** Figure 4**, Supp. Table 1).** We analyzed all G-R-F-x-V/I motifs identified due to the abundance of known motifs that contain a glycine a the -2 position. For the Δ_gly_-R-F-x-V/I and R-x-F-x-V/I we analyzed the top 20% of each group. 872 total motifs were analyzed **(Figure 3C)**.

We assigned motifs a score on a 100 point scale; with 100 being the best possible score **(Supp.** Figure 4**, Supp. Table 1)**. 10% of the score was assigned using NCBI BLAST to analyze conservation (Altschul et al., 1990). The motif had to be at least conserved in mouse (*Mus musculus*) or it was discarded, but it was scored higher if it was also conserved in frog (*Xenopus laevis*). 20% of the score was based on cellular localization. Since SPAK and OSR1 are present in the cytosol and nucleus, the motif had to be listed in the UniProt GO annotation as present in the cytosol, nucleus, or cytosol-facing region of membrane protein or it was discarded, and was scored lower if the motif was partly embedded in the membrane (Uniprot-Consortium, 2015). The largest part of the score (50%) was assigned based on solvent accessibility. The CCT domain must be able to access the motif without steric hinderance; thus, we took into account the degree of solvent accessibility utilizing primarily AlphaFold2 structural predictions or experimentally determined structures of identical or related proteins when no AlphaFold2 model was available (Jumper et al., 2021; Varadi et al., 2021). Motifs in loops and β-hairpins (2 strands) were scored the highest, followed by terminal β-strands of a sheet (≥ 3 strands), then internal β-strands and α-helices. Motifs also received a significantly lower score if they were packed against other parts of the structure, but a slightly higher score if the motif was partially a loop. For all three of the scoring metrics, if the information was unknown an intermediate score was assigned. Finally, we also took into account the initial Scansite score, which is based on motif sequence. Scansite scores range from zero to one with a lower score being better. Therefore 20% of the score was determined by the formula “(1-Scansite score) x 20” **(Supp. Figure 4, Supp. Table 1)**.

Of the 872 motifs analyzed 371 were discarded due to lack of conservation or cellular localization **(Figure 3C)**. The remaining 501 motifs were scored and ranked (higher score = better rank). 24 of the 27 previously validated motifs were present within the top 78 ranked motifs (top 2.1% of all 3675 motifs), and the majority of those clustered at the very highest rankings; providing compelling evidence that our scoring method worked exceptionally well **(Figure 3D,E)**.

Our results also suggest the presence of many additional uncharacterized SPAK/OSR1 CCT binding motifs since the majority of motifs clustered with the previously validated motifs have yet to be described **(Figure 3D,E)**. Gene ontology (GO) analysis of the top 50 genes associated with the highest ranked motifs indicate a range of biological processes **(Figure 3F, Supp. Table 2)** (Mi et al., 2013; Thomas et al., 2022). Some, such as protein phosphorylation, cell volume, and ion homeostasis, are expected and further serve as validation of the scoring method used for the predictions. Other biological processes are either not well described or completely lacking from WNK literature, such as DNA-templated transcription initiation, protein ubiquitination, p53 signaling, and plasma membrane protein localization. Therefore, our results imply there are rich, underexplored areas of biology associated with WNK signaling.

### Validation of a subset of predicted interacting proteins

We validated subsets of interactions biophysically by fluorescence anisotropy and in mammalian cell lysates by co-immunoprecipitation **(Figure 4,5)**. We selected a small group of proteins for this analysis either because of intriguing data in the literature or due to our preliminary data, noted below, that suggested potential relationships to WNK signaling. Several were a focus because of their connections, like WNK1, to blood pressure regulation. These include NOS3 (eNOS, endothelial nitric oxide synthase), a central regulator of the cardiovascular system and blood pressure; decreased NOS3 activity has been linked to increased risk of developing hypertension and a variety of other pathologies (Tran et al., 2022). The G protein-coupled receptor kinase 4 (GRK4) regulates sodium handling in the kidney via dopaminergic and renin–angiotensin signaling pathways, and mutations in the protein have been linked to hypertension (Jose et al., 2010; Villar et al., 2009; Wang et al., 2016). The K^+^-Cl^-^ cotransporter 4 (KCC4) is a well-documented substrate of OSR1/SPAK but to our knowledge the motif variant R-x-F-x-V had not been implicated in the interaction of the proteins (de Los Heros et al., 2014; Garzon-Muvdi et al., 2007). A family member (TSC22D3) of the protein TSC22 domain family protein 1 (TSC22D1) has been shown in a mouse knockout model to regulate Na^+^/K^+^ balance in the kidney, and WNK1 expression (Fiol et al., 2007; Rashmi et al., 2017). WNK1 also has well documented actions on protein localization and two of the predicted interactors were selected due to their known actions on protein trafficking. TBC1 domain family member 4 (TBC1D4) participates in the regulation of surface expression of glucose transporters, and WNK1 has been shown to regulate the function of TBC1D4 (Mendes et al., 2010). Cavin1 is an essential component of caveolae, which are important mediators of protein localization. Other potential interactors tested include Rho guanine nucleotide exchange factor 5 (ARHGEF5), like WNK1, often involved in the regulation of cell morphology and the actin cytoskeleton (Debily et al., 2004) and the autophagy-related protein 9A (ATG9A), an essential component of the autophagy machinery, because we previously showed that WNK1 inhibits autophagy (Gallolu Kankanamalage et al., 2016). Glycogen phosphorylase, liver form (PYGL), catalyzes the cleavage of glycogen into glucose monomers, which could be a means to regulate cellular osmolarity. Known interactors WNK1 and NKCC2 (Na^+^-K^+^-2Cl^-^ cotransporter) were used as controls.

**Figure 4.**
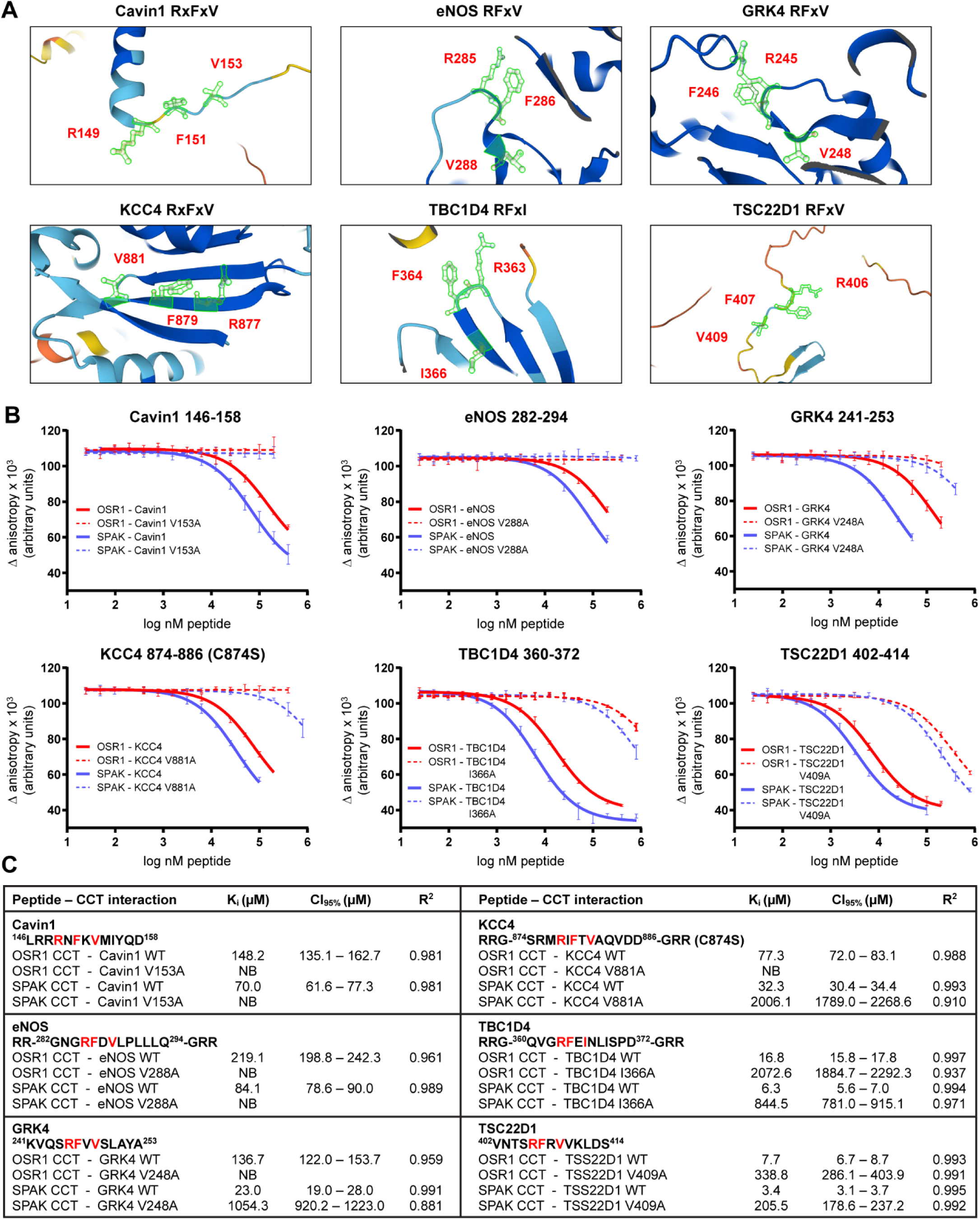
Multiple identified motifs interact with SPAK and OSR1 in vitro. **(A)** AlphaFold structural predictions of motifs in the proteins (Jumper et al., 2021; Varadi et al., 2021). Images generated using AlphaFold 3D Viewer. Green sidechains with red lettering indicate core motifs. Secondary structure coloring indicates model confidence. Dark blue: very high, light blue: high, yellow: low, orange: very low. Very low model confidence can indicate region is unstructured in isolation. Model confidence was not taken into account during structural analysis due to the high degree of complexity this would add to the scoring process across hundreds of structures. **(B)** Affinity determined by fluorescence anisotropy peptide competition assays. Unlabeled peptides containing predicted interaction motifs displace labeled peptide (NH_3_^+^-NLVGRF-[DAP-FAM]-VSPVPE-COO^−^] [diaminopropionic acid (DAP)). Labeled peptide held constant at 25 nM, SPAK CCT and OSR1 CCT are constant at 1.5 μM and 3.0 μM, respectively. OSR1 CCT with WT peptide: solid red line; OSR1 CCT with core motif mutation (V/I→A): dashed red line; SPAK CCT with WT peptide: solid blue line; SPAK CCT with core motif mutation (V/I→A): dashed blue line. **(C)** Quantitation of results in (B). Affinity reported as inhibition dissociation constant (K_i_, µM). Upper and lower limits of 95% confidence interval indicated. Fold-change decrease in affinity (fold-change increase in K_i_) relative to wild type indicated. Goodness of fit reported as R-squared (R^2^). All measurements are n=3. GraphPad Prism 10 with one site fit to K_i_ model used for data analysis. Red letters indicate core motif.

**Figure 5.**
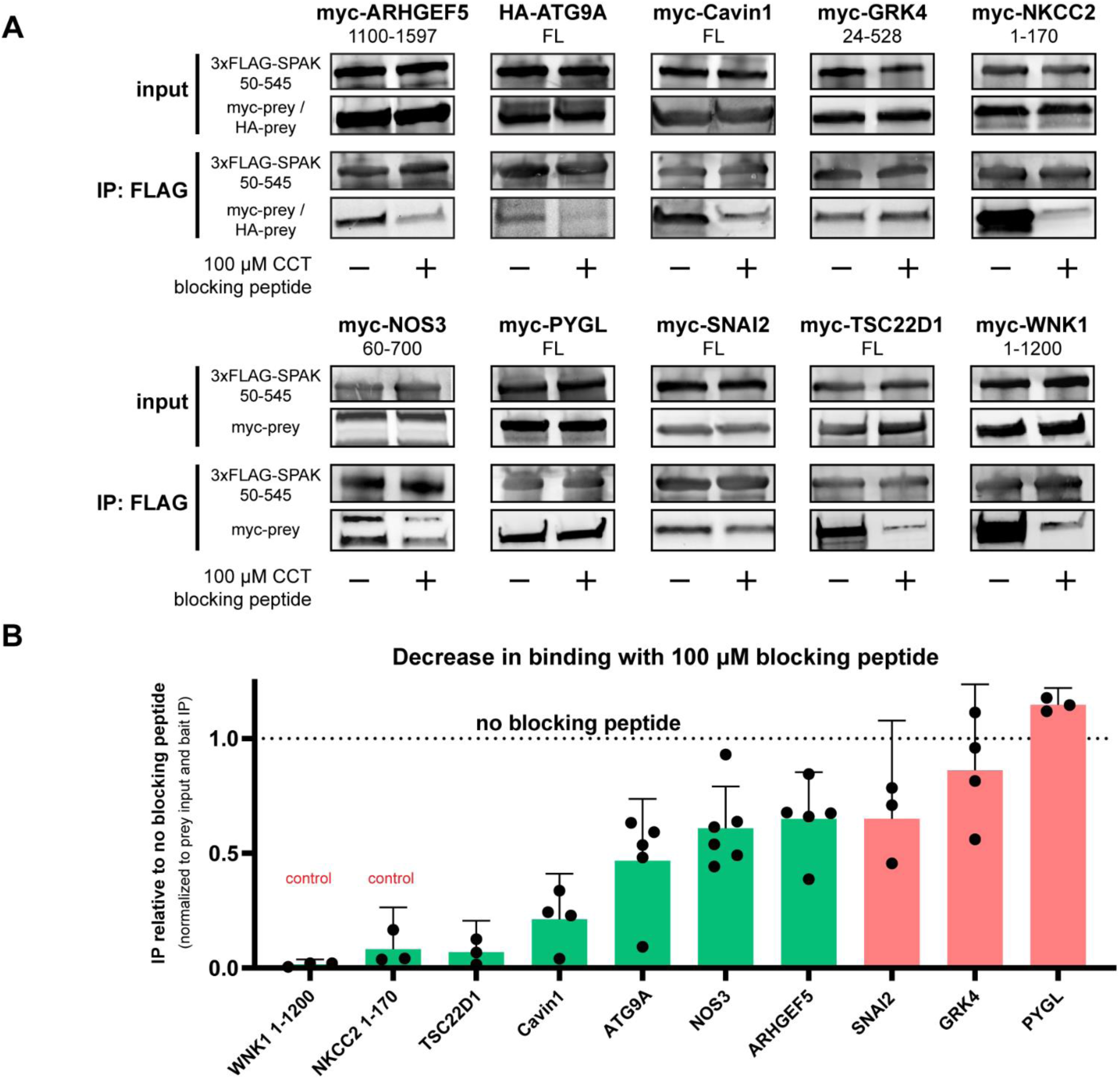
Validation of interactions via co-IP in HEK293T cells. **(A)** Overexpressed FLAG-hSPAK 50-545 bait protein was used to pull-down overexpressed myc-tagged prey proteins (HA tag for ATG9A) from HEK293T cell lysates. 100 µM CCT blocking peptide (WNK1 1253-1265; NH_3_^+^-SAGRRFIVSPVPE-COO^-^) was used as a control to block interactions. **(B)** Quantification of blots in (A). Error bars indicate 95% confidence intervals and replicates are shown as dots. Green are statistically significant and red are not.

13-mer peptides containing motifs of interest were interrogated by fluorescence anisotropy peptide competition assays identical to those described earlier **(Figure 1F)**. Here we show the results for the peptides that were soluble, and therefore compatible with our assay (Cavin1, KCC4, NOS3, TBC1D4, GRK4, and TSC22D1). Utilizing the AlphaFold2 structural models we found that four of the motifs were found in completely solvent-exposed loops, one was found in a partially exposed loop, and only KCC4 was found to be sandwiched in the middle of a β-sheet **(Figure 4A)**.

As mentioned, some of the peptides corresponding to the motifs in the other proteins we chose to investigate by co-immunoprecipitation were not compatible due to poor solubility, even with the addition of flanking arginines to enhance solubility. It should be noted that the introduction of flanking arginines to the peptides for KCC4, NOS3, and TBC1D4 to enhance solubility could alter the affinity of the peptides, but our primary goal was to demonstrate that mutation of the V/I in the core motif severely disrupts the interactions. Mutation of the V/I to alanine in the core motifs for all six proteins tested and for both SPAK and OSR1 CCT domains led to 1-2 orders of magnitude loss in affinity or no detectable binding, strongly suggesting these peptides are binding specifically to the CCT domains through engagement of the core motif **(Figure 4B,C)**.

We validated interactions by co-immunoprecipitation (co-IP) in HEK293T cell lysates. Anti-FLAG magnetic beads pre-loaded with 3xFLAG-SPAK 50-545 (human, lacking proline-alanine rich region) expressed in HEK293T cells was used to co-IP myc (or HA) tagged proteins (full length or fragments) that were overexpressed in HEK293T cells as well. As a control we used 100 µM CCT domain blocking peptide, which corresponds to the wild-type WNK1 sequence SAGRRFIVSPVE, to compete with protein specifically bound via the CCT domain interaction **(Figure 5)**. WNK1 and NKCC2 fragments were used as positive controls. The majority of the proteins tested showed a statistically significant reduction of binding in the presence of the blocking peptide (TSC22D1, Cavin1, ATG9A, NOS3, ARHGEF5). Two did not (GRK4, PYGL), although a lack of co-IP difference between control and blocking peptide treated cells does not prove that no interaction occurs; only that we failed to demonstrate that it occurs through the CCT in a statistically significant manner. We also included SNAI2 (aka transcription factor Slug) even though it was not ranked due to the position of the motif phenylalanine at the very N-terminus of the protein, and although we did observe a decrease in interaction with blocking peptide it was not statistically significant **(Figure 5)**. Based on results from this study, we also recently published the validated interactions of OSR1 with both TBC1D4 and sortilin (ranked 32) in another study (Jaykumar et al., 2024). Because the majority of the interactions tested met criteria for CCT domain binding, our methodology to predict interactions between CCT domains and motifs is capable of revealing connections between WNK signaling and previously unidentified proteins.

### WNK signaling influences Cavin1 phosphorylation

Cavin1, ranked fourth in our prediction analysis, is an essential component of caveolae, which are important mediators of localization of proteins such as transporters, channels, and other membrane proteins **(Table 1)** (Hill et al., 2008; Lamaze et al., 2017; Parton, 2018; Parton and del Pozo, 2013). In our co-immunoprecipitation validation of interaction partners Cavin1 showed both a strong interaction and decrease in interaction following addition of CCT blocking peptide **(Figure 5)**. Cavin1 is phosphorylated on Tyr156 following insulin or EGF (epidermal growth factor) stimulation, which is the second most commonly found Cavin1 post-translational modification (PhosphoSitePlus) and is a few residues away from the core R-x-F-x-V binding motif (Guha et al., 2008; Hornbeck et al., 2015; Humphrey et al., 2013; Liu and Pilch, 2016; Pilch and Liu, 2011). Inhibition of WNK signaling by both WNK1 siRNA-mediated knockdown and inhibition with the pan-WNK inhibitor WNK463 significantly decreases Tyr156 phosphorylation in human dermal microvascular endothelial cells (HDMEC) with little change in total Cavin1 protein amount **(Figure 6 A-D)**.

**Figure 6.**
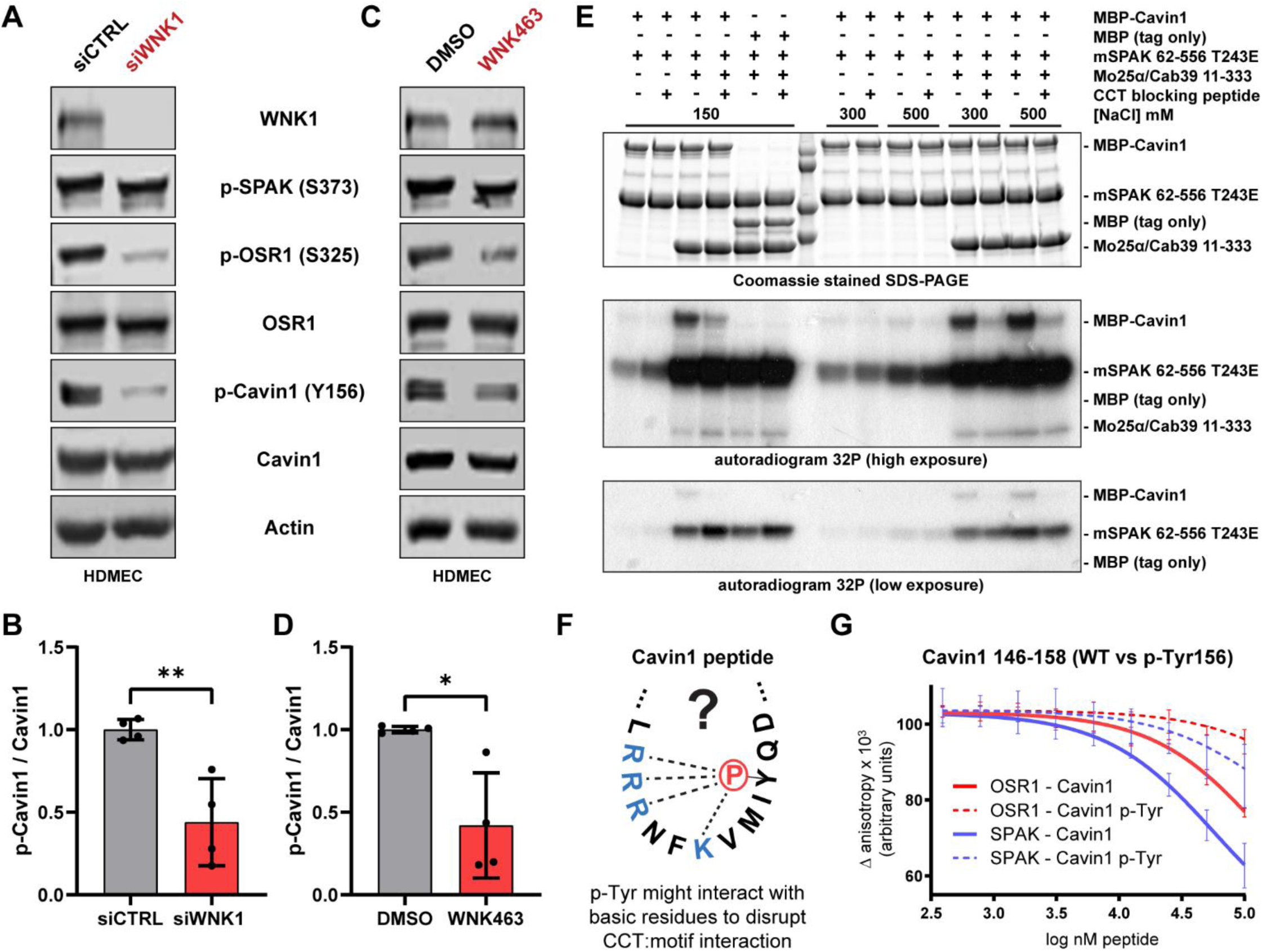
Cavin1 phosphorylation is influenced by WNK signaling. **(A-D)** siRNA-mediated knockdown of WNK1 (72 hours) and inhibition with the pan-WNK inhibitor, WNK463 (18 hours, 1 µM), in human dermal microvascular endothelial cells (HDMEC). Data reported as mean ± SD (n=3). **(E)** 32P incorporation into MBP-Cavin1. MBP refers to affinity tag maltose binding protein. Following 15 min preincubation, ATP was added and Cavin1 (5 µM) phosphorylation was monitored in the presence of constitutively active SPAK 62-556 T243E (15 µM) and the SPAK allosteric activating protein Mo25/Cab39 (15 µM). 100 µM CCT blocking peptide (WNK1 1253-1265; NH_3_^+^-SAGRRFIVSPVPE-COO^-^) was used to block the interaction between Cavin1 and SPAK. Phosphorylation reaction was run at varying NaCl concentrations for 10 minutes at 25 °C. Top panel is Coomassie stained gel and bottom two panels are 32P autoradiograms at different exposure times. MBP tag alone was used as a control (n=3). **(F)** Diagram indicating potential intramolecular interactions (dashed lines) between the acidic phospho-tyrosine 156 at the +5 position of the Cavin1 motif and multiple basic residues at the -4, -3, -2, and +1 positions. Such an interaction could potentially diminish the ability of CCT domains to interact with the motif. **(G)** Affinity determined by fluorescence anisotropy peptide competition assays. Unlabeled peptides were used to displace labeled peptide (NH_3_^+^-NLVGRF-[DAP-FAM]-VSPVPE-COO^−^] [diaminopropionic acid (DAP)). Labeled peptide held constant at 25 nM, SPAK CCT and OSR1 CCT are constant at 1.5 μM and 3.0 μM, respectively. OSR1 CCT with Cavin1 peptide lacking p-Tyr156: solid red line; OSR1 CCT with p-Tyr156 Cavin1 peptide: dashed red line; SPAK CCT with Cavin1 peptide lacking p-Tyr156: solid blue line; SPAK CCT with p-Tyr156 Cavin1 peptide: dashed blue line. Higher line indicates weaker binding. Quantification unavailable due to instability of Cavin1 p-Tyr peptide at higher concentrations (n=3).

In addition to Tyr156 phosphorylation, in vitro utilizing purified proteins, constitutively active mouse SPAK 62-556 T243E phosphorylated MBP-Cavin1 (MBP: maltose binding protein), no phosphorylation of the MBP tag alone was observed, and addition of the CCT blocking peptide dramatically decreased phosphorylation **(Figure 6E)**. Addition of the SPAK/OSR1 kinase activity-enhancing protein Mo25α (aka Cab39) also significantly increased phosphorylation, as has been shown for other substrates (Filippi et al., 2011). Higher NaCl concentrations modestly increased Cavin1 phosphorylation.

Finally, we postulated that because there are several basic residues close to the core binding motif including a triple arginine (RRR) motif that partially comprises the core motif, perhaps phosphorylated Tyr156 forms an electrostatic interaction with these residues and occludes CCT domain binding to the motif **(Figure 6F)**. In our FP-based peptide competition assay the motif peptide (Cavin1 146-158) exhibited a decrease in binding affinity to both the SPAK and OSR1 CCT domains when Tyr156 was phosphorylated **(Figure 6G)**.

While detailed mechanisms are missing, our findings implicate WNK signaling in aspects of Cavin1 function, SPAK/OSR1 as direct interactors of Cavin1, and suggest a possible mechanism for regulation of the interaction between the kinases and Cavin1 via phosphorylation at Tyr156.

### TSC22D1 interacts with SPAK/OSR1 and influences WNK pathway signaling

Several lines of evidence point to a connection between TSC22D1 and WNK signaling. TSC22D proteins were originally identified as TGF-β regulated transcription factors, and we have previously shown a connection between them and WNK and TGF-β-mediated signaling (Jaykumar et al., 2022; Lee et al., 2007; Shibanuma et al., 1992). Three of the four TSC22D family members contain strongly-predicted binding motifs with ranks of 3, 8, and 77 for TSC22D1, TSC22D4, and TSC22D2, respectively **(Table 1**, **Figure 7A)** and all four can heterodimerize through their conserved leucine zippers (Kester et al., 1999). Similar to Cavin1, TSC22D1 was one of the strongest candidates for further analysis based on our co-immunoprecipitation validation assays **(Figure 5)**. We found that endogenous TSC22D1 also co-immunoprecipitated with 3xFLAG-OSR1 and the CCT blocking peptide prevented their association in HEK293T cells (**Figure 7B)**. Endogenous TSC22D1 is also pulled down with GST-OSR1 CCT but not GST alone **(Figure 7C)**.

**Figure 7.**
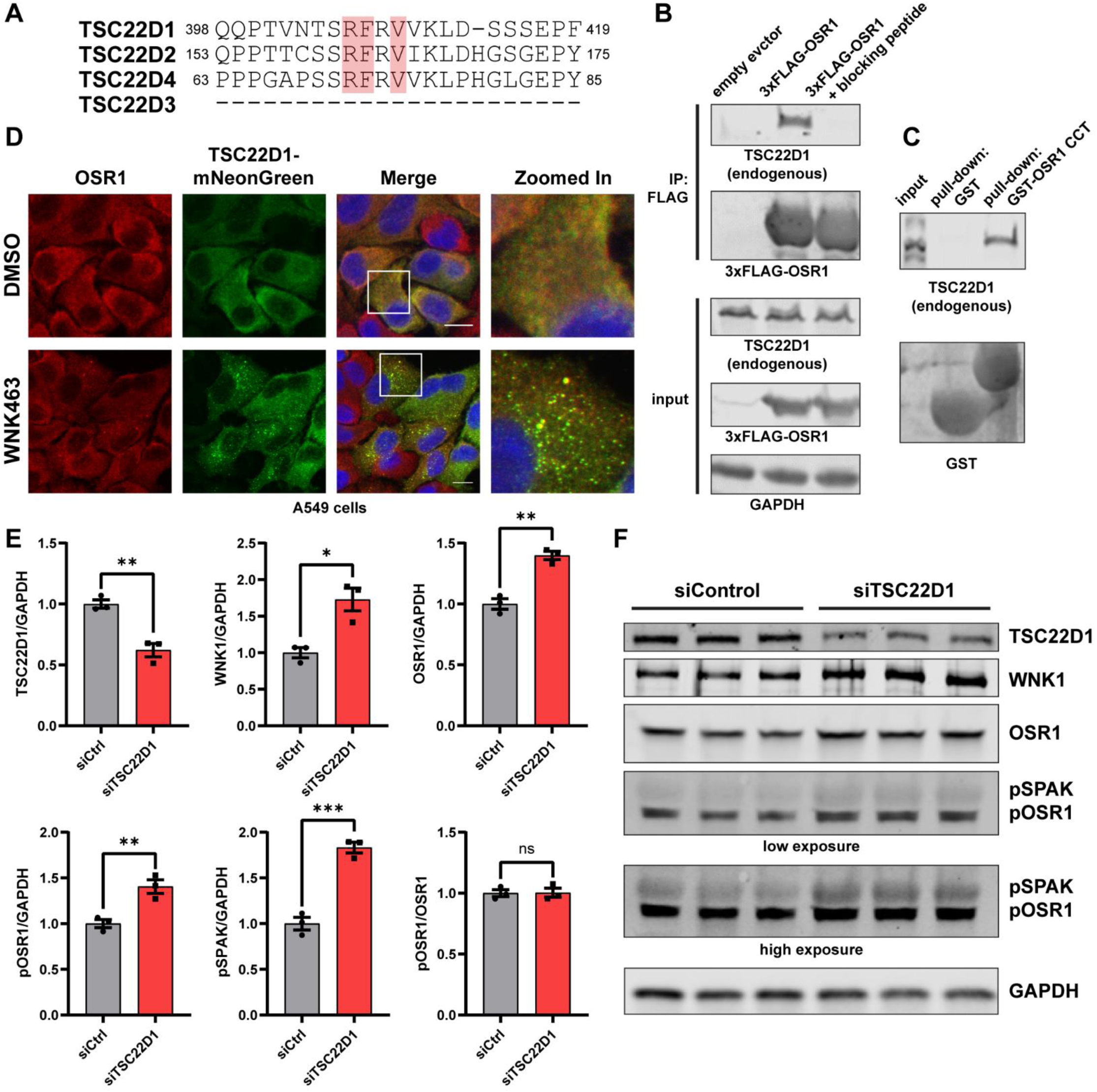
TSC22D1 interacts with SPAK/OSR1 and impacts WNK signaling. **(A)** Multiple sequence alignment of TSC22D proteins around identified R-F-x-V motif. TSC22D3 lacks the motif. **(B)** co-IP of endogenous TSC22D1 with 3xFLAG-OSR1 in HEK293T cells. 50 µM CCT blocking peptide (WNK1 1253-1265; NH_3_^+^-SAGRRFIVSPVPE-COO^-^) was used as a control to inhibit the interaction (n=3). **(C)** Similar to (B) except bacterially expressed GST-tagged OSR1 CCT domain was used to pull-down TSC22D1 from HEK293T cell lysates (n=3). **(D)** Colocalization of endogenous OSR1 and stably-expressed TSC22D1-mNeonGreen in A549 cells in the presence or absence of 1 µM pan-WNK inhibitor, WNK463, treated 24 hours (n=3). **(E and F)** Western blot of partial siRNA-mediated knockdown of TSC22D1 in HDMEC after 48 hours siRNA treatment. Data are reported as mean ± SEM (n=3). pSPAK/pOSR1 blot shown twice at two different intensities.

We were unable to validate the TSC22D1 antibodies for immunolocalization, even though they worked well for immunoblotting. As a result, we tagged TSC22D1 with mNeonGreen and made stable A549 lung cancer and HEK293T cell lines expressing the tagged protein (TSC22D1-mNeonGreen). Treatment with the pan-WNK inhibitor, WNK463 (1 µM, 4 hours), induced TSC22D1 puncta in both cell types **(Figure 7D, Supp.** Figure 5A). Puncta were smaller and more evenly distributed in A549 cells compared to HEK293T cells. In A549 cells endogenous OSR1 colocalized in puncta with tagged TSC22D1 in the presence of WNK463 **(Figure 7D)**. In HEK293T cells, treatment with hypotonic media also led to a decrease in TSC22D1 puncta size **(Supp. Figure 5A)**.

Partial siRNA-mediated knockdown of TSC22D1 significantly altered amounts of WNK1 and OSR1 in HDMEC **(Figure 7 E,F)**. OSR1 and SPAK activation were also significantly increased, but the increase in phosphorylation is likely related to increased protein expression as there was no change when pOSR1 was normalized to total OSR1 **(Figure 7E)**. Treatment of HDMEC with WNK463 also altered mRNA amounts of TSC22D family members **(Supp. Figure 5B)**. Taken together these findings point to SPAK and OSR1 directly interacting with TSC22D family members, and a connection between the expression of TSC22D family members and components of the WNK pathway.

### CCT-like domains and alternative binding motifs utilize a common binding mode

Each of the four WNKs contain two CCT-like (CCTL) domains (CCTL1 and CCTL2), as was initially reported for WNK1 and later other WNK family members (Bartual et al., 2017; Moon et al., 2013; Pinkas et al., 2017; Taylor and Cobb, 2022). Structural predictions provided by AlphaFold2 indicate all WNK CCTL domains share a common structure and motif binding pocket with SPAK and OSR1 CCT domains (Jumper et al., 2021; Varadi et al., 2021). Using AlphaFold2, we also found that NRBP1 and NRBP2, which are the closest relatives to WNKs in the human kinome tree, also contain a CCTL domain **(Figure 8A,B)** (Manning et al., 2002). WNK1 CCTL1 and WNK2 CCTL1 have been shown to interact with motif-containing peptides in an in vitro setting using purified proteins, although no physiological evidence for a function of motif:CCTL interactions has been demonstrated (Moon et al., 2013; Pinkas et al., 2017).

**Figure 8.**
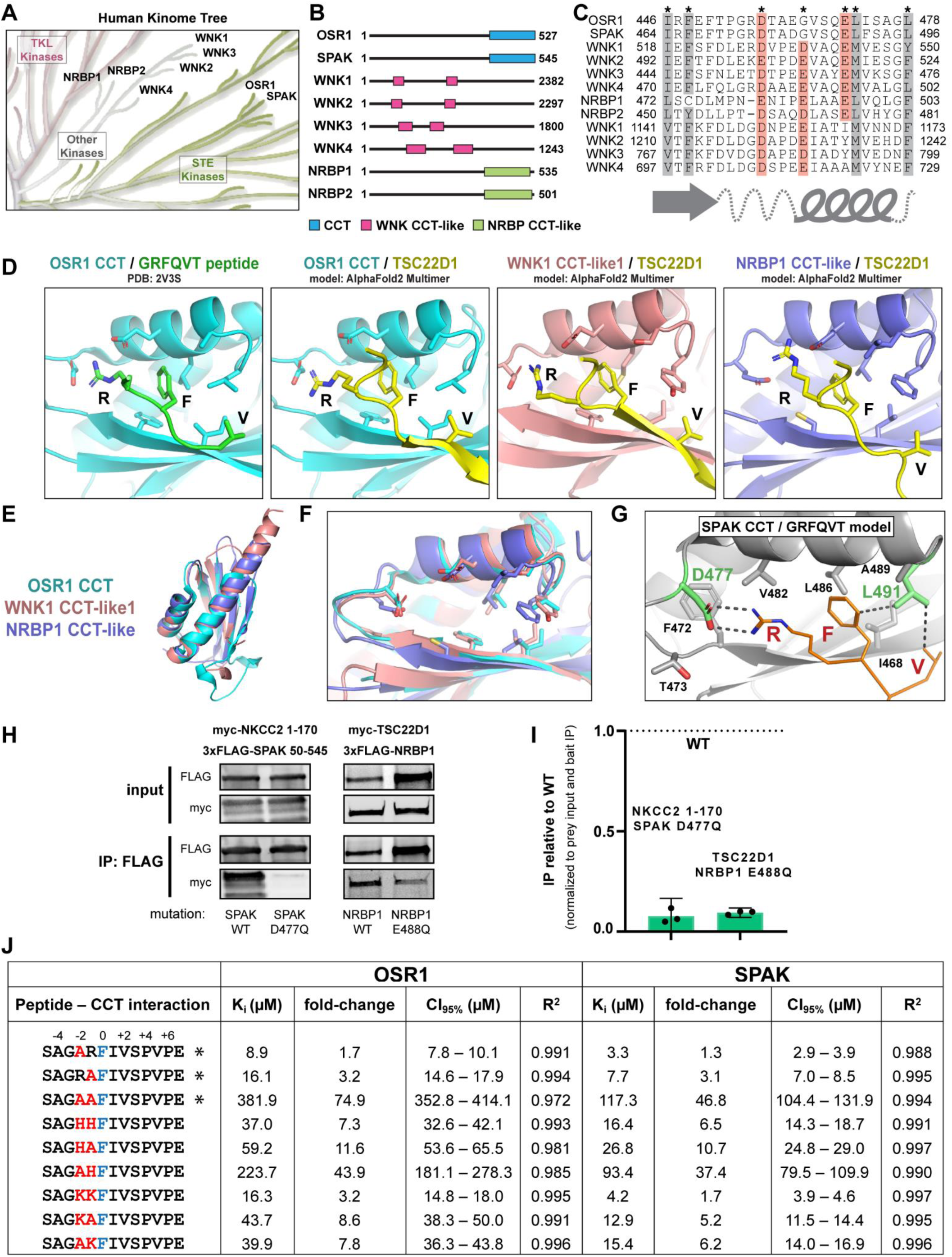
CCT-like domains and alternative binding motifs expand complexity. **(A)** Region of human kinome tree containing WNK1-4, SPAK/OSR1, and NRBP1/2; all proteins containing CCT or CCT-like (CCTL) domains (Manning et al., 2002). **(B)** Diagram depicting locations of CCT and CCTL domains in WNKs, NRBPs, and SPAK/OSR1. **(C)** Multiple sequence alignment of regions of CCT and CCTL domains that comprise the motif binding pocket, which consists of a β-srand – loop – α-helix. Conserved residues important for binding are indicated. **(D)** Comparison of the crystal structure of the OSR1 CCT bound to a GRFQVT hexapeptide (left panel, PDB:2V3S) with AlphaFold2 Multimer-derived models of OSR1 CCT, WNK1 CCTL1, and NRBP1 CCTL bound to the RFxV motif of TSC22D1 (Cianfrocco et al., 2017; Evans et al., 2022; Jumper et al., 2021; Varadi et al., 2021). **(E and F)** Structural alignment of AlphaFold2 models of CCT and CCTL domains in (D). Residues important for binding are depicted as sticks. **(G)** Model of SPAK bound to the GRFQVT peptide made by overlaying the OSR1 CCT:GRFQVT structure on the crystal structure of the SPAK CCT domain (PDB: 7O86). D477 (green) was mutated in the present study to prevent binding while L491 (green) was mutated in a previous study to prevent interactions (Zhang et al., 2015). **(H and I)** 3xFLAG-tagged SPAK 50-545 ± D477Q or NRBP1 ± E488Q were used to co-IP myc-NKCC2 1-170 and myc-TSC22D1, respectively, in HEK293T cells. Results are plotted as mean ± SEM of mutants relative to WT (n=3). **(J)** Affinity determined by fluorescence anisotropy peptide competition assays. Unlabeled peptides at various concentrations displace labeled peptide (NH_3+_-NLVGRF-[DAP-FAM]-VSPVPE-COO_−_] [diaminopropionic acid (DAP)). Labeled peptide held constant at 25 nM, SPAK CCT and OSR1 CCT are constant at 1.5 μM and 3.0 μM, respectively. Red letters are mutated relative to the wild type (WT) sequence. Blue letter is motif phenylalanine (position 0). Affinity reported as inhibition dissociation constant (K_i_, µM). Upper and lower limits of 95% confidence interval indicated. Fold-change decrease in affinity (fold-change increase in K_i_) relative to wild type indicated. Goodness of fit reported as R-squared (R_2_). All measurements are n=3. Fit curves and fluorescent probe affinity measurement can be found in **Supp. Fig. 3.** GraphPad Prism 10 with one site fit to K_i_ model used for data analysis. Asterisk (*) indicates results previously published (Taylor et al., 2018).

While there are slight differences in the topology of CCT and CCT-like domains, the overall folds are strikingly similar; particularly the core motif binding pocket composed of a β-strand followed by a loop and then an α-helix where all the hydrophobic and acidic residues that bind the core motif are located **(Figure 8 C-F)**. Utilizing AlphaFold2 Multimer we were able to reliably generate models of TSC22D1 R-F-x-V motif interactions with OSR1 CCT, WNK1 CCTL1, and NRBP1 CCTL **(Figure 8D)** (Cianfrocco et al., 2017; Evans et al., 2022; Jumper et al., 2021). Previous work studying orthologs of TSC22D and NRBP proteins in Drosophila identified regions within the proteins corresponding to the R-F-x-V motif and CCTL domain that were able to co-immunoprecipitate (Gluderer et al., 2010). Additionally, TSC22D2, WNK1, and NRBP1 share the highest degree of codependence in CRISPR and RNAi screens according to DepMap.org (Tsherniak et al., 2017). These findings suggested that NRBP CCTL domains might interact with TSC22D R-F-x-V motifs **(Figure 7A)** in a manner similar to SPAK/OSR1 CCT domains.

We identified a point mutation in the SPAK CCT domain that significantly diminished motif binding. Previous work in mice had identified a point mutation (mouse L502A, human L491A) that drastically reduced motif interactions **(Figure 8G)** (Zhang et al., 2015). However, because this mutant also impacts hydrophobic interactions required for CCT domain folding we chose to pursue an alternative mutant, D477Q (human). This point mutant neutralizes a charge and increases side chain length on a residue critical for interactions with the motif arginine **(Figure 8G)** (Elvers et al., 2022; Villa et al., 2007). We tested several mutants by FP with purified SPAK CCT domain, the D477Q mutant led to an approximately 50-fold decrease in peptide probe binding affinity **(Supp. Figure 6A)**. Interestingly, D477A had been previously shown to diminish interactions with WNK1 and NKCC1 (Vitari et al., 2006). By co-immunoprecipitation of myc-NKCC2 1-170 with 3xFLAG SPAK 50-545 in HEK293T cells, the mutant led to a near complete loss of interaction **(Figure 8H,I)**. A similar result was observed for Cavin1 **(Supp. Figure 6B)**. We then utilized this same strategy in the NRBP1 CCTL domain. NRBP1 E488Q also neutralizes the charge in the position analogous to D477, although it does not alter side chain length. As expected, E488Q diminished the interaction between myc-TSC22D1 and 3xFLAG-NRBP1, suggesting that NRBP1 interacts with the TSC22D1 R-F-x-V motif in a manner similar to SPAK/OSR1.

Beyond alternative CCTL domains, we also investigated whether alternative core binding motif sequences might be able to interact with CCT domains. We previously reported that mutation of arginine to alanine in the WNK1 peptide sequence (SAGRRFIVSPVPE) at either the -3 or -2 positions both still bound to CCT domains with only a modest decrease in affinity illustrating that R-x-F-x-V/I motifs were viable core motifs (Taylor et al., 2018). We analyzed these data again with current software and found similar results **(Figure 8J; Supp. Figure 3)**. Alanine substitution at either the -3 or -2 positions led to an approximate 1.5-fold or 3.1 fold drop in affinity respectively, whereas mutation of both leads to a drastic decrease in affinity.

We then analyzed effects of substitution of the arginines in the WNK1-derived peptide to other basic residues **(Figure 8J; Supp. Figure 3)**. Replacing the arginines with histidine-histidine led to an approximate 7-fold decrease in affinity, but the affinities were still in the low-mid micromolar range, which could provide enough specificity for signaling, and the same is true for histidine-alanine (11-fold decrease), but the decrease for alanine-histidine was more substantial leading to K_i_ values in the high micromolar range. Surprisingly, replacement with lysine-lysine led to K_i_ of 16.3 µM and 4.2 for OSR1 and SPAK, respectively, which is only a slight loss in affinity compared to wild type, and while both lysine-alanine and alanine-lysine led to a further decrease in binding, all the measured affinities were still in the low-mid micromolar range.

Our results combined with the results of previous studies and structural predictions indicate a high degree of complexity in how CCT and CCT-like domains contribute to signaling both within and outside the WNK pathway. More than one CCT or CCT-like domain may interact with the same motifs with varying degrees of specificity providing an additional layer of complexity to signaling by both providing a means of motif competition but also by bringing different signaling proteins together such as in the case of motif-containing proteins that oligomerize, notably WNKs, TSC22Ds, and Cavin1. Additionally, alternative motifs outside of the classic R-F-x-V/I and more recently identified R-x-F-x-V/I may also function to facilitate viable protein:protein interactions both within and outside the WNK pathway.

## DISCUSSION

Protein-protein interactions facilitate the formation of short- and long-lived protein complexes designed to communicate with cellular processes with specificity. Outside their kinase domains, WNKs are currently viewed as largely unstructured and function as scaffolds through several sequence and structural properties, including Pro-rich regions, coiled-coils and R-F-x-V motifs that mediate interactions with the OSR1 and SPAK CCT domains. WNKs activate OSR1 and SPAK by phosphorylation, enabling OSR1 and SPAK to phosphorylate a variety of substrates. The capacity of the CCT domains to recognize many protein targets through these common short motifs enables inputs from OSR1/SPAK and WNKs to extend to many cellular regulatory pathways.

More conventional screening methods used to identify direct protein:protein interactions such as affinity-purification mass spectrometry (AP-MS), yeast two-hybrid, and BioID are powerful, but often limited by the inclusion of false-positives (nonspecifically-bound proteins, indirect interactions, irrelevant proteins in close spatial proximity) and false-negatives (weak interactions, low expression). This study was designed to minimize these limitations to identify direct protein-protein interactions that underlie processes related to WNK pathway function, and to accelerate characterization of the essential biochemical mechanisms that encompass both dominant and more modest regulatory control that will help describe a bigger picture of WNK signaling.

Protein phosphorylation/dephosphorylation signaling pathways often depend on regulated interactions to decide pathway specificity and function by identifying context-specific targets, feedback mechanisms, and localization. Phosphoprotein phosphatases 1 and 2A (PP1 and PP2A) acquire specificity through targeting subunits that recognize and restrict substrates. Interestingly, PP1 also uses a short linear interaction motif to identify regulatory subunits and substrates, including the OSR1/SPAK substrate, the Na^+^, K^+^ 2Cl^-^ cotransporter (Darman et al., 2001; Gagnon and Delpire, 2010). PP1 substrate recognition is an example of how this common type of short motif permits incredibly broad control of many cellular processes.

Among highly predicted CCT binding partners, no mechanistic basis for TSC22D protein interactions with the WNK pathway was previously known. Originally identified as TGF-β regulated transcription factors, TSC22D proteins are intertwined with the WNK pathway from proteomics and other studies, with little biochemical insight. The four TSC22D proteins heterodimerize and TSC22D1, 2, and 4 contain binding motifs that are highly ranked. We show here that TSC22D1 can co-localize with OSR1 and its localization is sensitive to WNK inhibition. In a mouse knockout model, the Pearce lab showed that TSC22D3 (GILZ) regulates Na^+^/K^+^ balance in the kidney, and has a strong effect on WNK1 expression (Rashmi et al., 2017). Transcription of TSC22D family members increases following hypertonic stress (Fiol et al., 2007). TSC22D2 and TSC22D4 interact with SPAK as well as WNKs as deduced from large-scale mass spectrometry-based proteomics studies (BioGRID database) (Oughtred et al., 2021) and TSC22D2 was shown to interact with SPAK and WNK3 in a yeast two-hybrid screen (Li et al., 2016). Surprisingly, TSC22D2 is also the molecule with results most similar to WNK1 in analysis of siRNA and CRISPR screens from data aggregated in Depmap.org. (Tsherniak et al., 2017). Knowing how they bind should assist in determining more about their inter-related functions.

A second highly predicted and validated binder, Cavin1 is a compelling target for future investigation because its mechanosensitivity in the endothelium may be related to WNK1 functions in endothelial migration, TGF-β sensitivity, and responses to mechanical stress [Dbouk et al, Jaykumar et al, 2022; Jung et al, in preparation]. Cavin1, ranked fourth in our prediction analysis, is an essential component of caveolae, which are important mediators of localization of proteins such as transporters, channels, and other membrane proteins **(Table 1)** (Hill et al., 2008; Lamaze et al., 2017; Parton, 2018; Parton and del Pozo, 2013). Cavin1, also referred to as PTRF (Polymerase I and transcript release factor), is required for efficient ribosomal biogenesis by dissociating paused RNA polymerase I from transcriptional complexes (Jansa and Grummt, 1999; Liu and Pilch, 2016). In addition to its identification as a pathway target through OSR1/SPAK binding, our study shows the parallel control of tyrosine phosphorylation of Cavin1 by WNK1. Phosphorylation of Tyr156 has been implicated in ribosomal RNA transcriptional regulation, as a Tyr to Phe mutation reduces ribosomal RNA transcription (Jansa and Grummt, 1999; Liu and Pilch, 2016). Phosphorylation of Tyr156 can be catalyzed by receptors for epidermal growth factor (EGF) and insulin. In previous work we showed that depletion of WNK1 results in increased degradation of the EGF receptor (Jung and Cobb, 2023). Among possible explanations for the decrease in Cavin1 pY156 occurring as a consequence of WNK1 loss, is a reduction in plasma membrane EGF receptor needed to catalyze the reaction.

In a number of cases, phosphorylatable residues lie in close proximity to CCT binding motifs. As noted in **Figure 2B**, more than half of the already published motifs are directly followed at the +3 position by Ser or Thr. Work initially by the Alessi/van Aalten labs using short peptides suggested that CCT domain-motif binding is regulated by phosphorylation of such residues (Villa et al., 2007). Although less dramatic, we also see differences in peptide binding as a result of Ser phosphorylation of a WNK1 peptide, as well as phosphorylation of Tyr at the +5 position in a Cavin1 peptide. The common occurrence of Ser/Thr/Tyr near motifs indicates that phosphorylation may be a frequent mechanism controlling CCT domain-target binding.

Some of the proteins found to interact link the WNK1 pathway mechanistically to previously identified functions; these include sortilin and TBC1D4. We tested TBC1D4, ranked at 246, because of previous work from the Jordan lab indicating an association with WNK1 and glucose transporter trafficking (Henriques et al., 2020; Mendes et al., 2010) and because of earlier studies suggesting effects of WNK1 on glucose homeostasis (Nishida et al., 2012). In recent work, we found that TBC1D4, also known as AS160, and a second predicted OSR1/SPAK interactor sortilin, ranked at 28, cooperate to maintain insulin-dependent Glut4 trafficking, uncovering one mechanism using two CCT interactions through which WNK1 can influence glucose metabolism (Jaykumar et al., 2024). In work not yet published, we were able to show that OSR1 co-immunoprecipitates with endogenous ATG9A (Kannangara, et al, in preparation), supporting the association observed in vitro here. ATG9A is a core autophagy protein that provides another connection to our earlier findings that WNK1 inhibits autophagy (Gallolu Kankanamalage et al., 2016).

In addition to testing highly predicted interactions, we examined binding determinants of the OSR1/SPAK CCTs in detail to attempt to define limits of CCT-core motif interactions. We also touched upon the relevance of such motifs to interactions with CCT-like domains in WNKs and NRBPs. What we found was that sequence differences in the core motif, for example, ones in which arginine is replaced by other basic residues (e.g., 140 KK motifs in human proteome), may lead to physiologically important motif variants with only minor decreases in affinity. We also found that sequences outside the core motif can significantly influence binding specificity to the extent that motifs that appear poor may work well because of residues beyond the core sequence. The potential multivalency of interactions may be the feature that most complicates what early on appeared to be a simple domain-motif association. More than one CCT or CCT-like domain may interact with the same motifs with varying degrees of specificity providing an additional layer of complexity to signaling by both providing a means of motif competition and also by bringing different signaling proteins together such as in the case of motif-containing proteins that oligomerize, e.g., TSC22D proteins and Cavin1, and proteins with CCT-like domains that themselves oligomerize. WNKs oligomerize, as do OSR1 and SPAK, and also use phase separation to collaborate with other proteins (Boyd-Shiwarski et al., 2022; Boyd-Shiwarski et al., 2017; Kester et al., 1999; Kovtun et al., 2014; Lee et al., 2009; Lenertz et al., 2005; Taylor et al., 2015; Villa et al., 2008). We conclude that there is a high degree of complexity in how CCT and CCT-like domains contribute to signaling both within and outside the WNK pathway that far exceed a simple 1:1 relationship.

## METHODS

### Protein purification and peptide production

His_6_-tagged CCT domains (mouse SPAK 459-556 and human OSR1 433-527) and mouse SPAK 62-556 T243E were expressed in Rosetta (DE3) E.coli (Novagen). Cells were grown at 37 °C, 225 rpm until OD_600_ reached 1.0 and then induced with 0.5 mM IPTG overnight at 20 °C, 150 rpm. Cells were lysed by freeze/thaw (-20 °C) and then with 1 mg/ml lysozyme in 50 mM Tris-HCl, pH 8.0, 300 mM NaCl, 10% glycerol, and protease inhibitors (1 mM phenylmethanesulfonyl fluoride, 583 µM pepstatin A, 762 µM leupeptin, 10.6 mM Nα-Tosyl-L-arginine methyl ester HCl, 10.8 mM tosyl-lysine-chloromethylketone HCl, 11.3 mM Nα-benzoyl-L-arginine methyl ester carbonate, 200 µM soybean trypsin inhibitor), on ice for 30 min followed by sonication. Soluble proteins were applied to nickel-nitrilotriacetic acid-agarose (Ni^2+^-NTA) and eluted with 500 mM imidazole gradient. Fractions analyzed by polyacrylamide gel electrophoresis in sodium dodecyl sulfate (SDS-PAGE), and dialyzed at 4 °C overnight. CCT domains dialyzed into 25 mM Tris-HCl pH 7.75, 125 mM NaCl, 1 mM DTT, and SPAK 62-556 T243E dialyzed into 20 mM HEPES pH 7.5 100 mM NaCl, 1 mM DTT. Proteins were subjected to Superdex200 gel filtration and fractions were pooled. GST-Mo25α 11-333 was purified similarly except 5 mM DTT and 1 mM EDTA were added to lysis buffer, glutathione beads were used for affinity purification followed by on-bead cleavage with TEV protease to remove tag and incubation with Ni-NTA beads to remove TEV, and the gel filtration buffer was 20 mM HEPES pH 7.5 100 mM NaCl, 1 mM DTT. MBP-Cavin1-His_6_ required special considerations for purification due to instability. Induction in Rosetta2 cells immediately arrests cell growth. Therefore, cells were grown to an OD_600_ of 2.0 prior to addition of 1 mM IPTG and growth overnight at 16 °C. Protein was first affinity purified similarly as His_6_-tagged proteins, then bound to amylose resin followed by on-bead cleavage in 20 mM HEPES pH 7.5, 500 mM NaCl, 1 mM DTT, and protease inhibitors/EDTA. Protein was subjected to Superdex 200 gel filtration.

### CCT domain specificity analysis by peptide array

Immobilized peptide membranes were purchased from Kinexus Bioinformatics Corp. Approximately 200 nmol of N-terminal acetylated peptides per spot was chemically synthesized on a TOTD membrane by coupling of the C-terminus of the peptide to the membrane and peptide synthesis was carried out on the membrane. Membranes were rinsed with EtOH then washed 30 mins with TBST (0.1 % Tween). Membranes were blocked 60 mins with 1:1 Odyssey blocking buffer to TBST (blocking solution). Membrane was then incubated with 300 µM His_6_-SPAK CCT (residues 449-545) or His_6_-OSR1 CCT (residues 433-527) in blocking solution for 60 mins. Membrane was then washed 2x5 mins in TBST. Membrane was then incubated with 1:1000 mouse anti-His6 antibody (Clontech) for 15 mins. Membrane was then washed 2x5 mins in TBST followed by incubation with 1:500 LI-COR IRDye 680RD mouse secondary antibody for 15 minutes. Membrane was then washed 4x5 mins in TBST before quantification by LI-COR Odyssey. Antibody-only overlay (background) spot intensities were subtracted from the CCT overlay spot intensities. The average spot intensity for each array was used to normalize intensity values between arrays.

### Peptide binding affinities by fluorescence polarization

1.5 µM His_6_-SPAK CCT or 3.0 µM His_6_-OSR1 CCT was combined with 25 nM NH_3_^+^-NLVGRF[DAP-FAM]VSPVPE-COO^-^ (DAP-FAM: 2,3-diaminopropionic acid, unnatural amino acid, conjugated to FAM) in a buffer of 25 mM Tris-HCl pH 7.75 (at 25° C), 125 mM NaCl, and 1 mM DTT. Unlabeled competing peptides were then added and subjected to a 2-fold stepwise dilution. Fluorescence polarization measurements were carried out on a BioTek Synergy H1 multi-mode plate reader equipped with a fluorescein polarizing filter cube (Em: 485 nm, Ex: 528 nm, 510 nm dichroic mirror). Data was fit to a model of one-site competitive binding to determine K_i_ using GraphPad Prism software. For measurements of binding affinity for labeled peptides to CCT domains a similar protocol was used but CCT concentration was varied. 1 mg/ml BSA was used for all peptides where the sequences were not derived from WNK1 or WNK4 in order to prevent nonspecific binding at higher competing peptide concentrations. All peptides initially diluted in 25 mM Tris-HCl pH 7.75 (at 25° C), 125 mM NaCl to 10 mM. DMSO added to peptides with solubility issues.

### Motif search, ranking, and analysis

Average spot intensities from the OSR1 CCT peptide arrays for residues outside the core R-F-x-V/I or R-x-F-x-V/I motifs were used as input for the Scansite3 webserver (scansite3.mit.edu) (Obenauer et al., 2003). Core motif phenylalanine was fixed in center of matrix. As input for positions corresponding to residues R and F in the core motif no other residues were allowed, and for the position corresponding to V/I the ratio of scores for V:I was 2 (based on difference in binding affinities determined by fluorescence polarization for FAM-QLVGRFQVTSSKE and FAM-QLVGRFQITSSKE). For the G-R-F-x-V/I motif class, G was also fixed, and rows 2, 3, 6, and 8-10 of the peptide array were used for scoring. For the Δ_gly_-R-F-x-V/I motif class, all residues except G were allowed at the position corresponding to Δ_gly_, and rows 1, 3, 6, and 8-10 were used for scoring. For the R-x-F-x-V/I motif class rows 6 and 8-12 were used for scoring. The UniProt *H. sapiens* database was searched for all valid motifs and the motifs were scored based on the inputs described above (Uniprot-Consortium, 2015). Scores were then used to rank motifs (lower score equates to higher probability of interaction).

Cutoffs for additional analysis were top 20% for Δ_gly_-R-F-x-V/I and R-x-F-x-V/I motifs. No cutoff was used for G-R-F-x-V/I motifs. Motifs not conserved in Mus musculus (mouse) were discarded. Motifs not annotated in UniProt GO terms as being present in either cytosol, nucleus, or within a membrane protein region within or partially in the cytosol were discarded (Uniprot-Consortium, 2015). Motifs containing a proline at the +1 or +3 positions were discarded. Motifs were scored according to **Supp. Figure 4.** AlphaFold2 was used when possible; otherwise the protein sequence was used for BLAST search of experimental-derived structures and the identical or closest-related protein was used by aligning motif to the sequence and determining the position in the structure. If a motif was present in a loop or β-hairpin that was over 100 residues away from surrounding structural elements it was packed/folded with it was treated as an isolated structural element. If any part of a motif was present in secondary structure or annotated as partial membrane helix it was treated as that element but assigned partial loop. After total analysis all three motif classes were combined and ranked by the overall score based on the 100 point system described in **Supp. Figure 4**.

Gene ontology analysis was performed using the top 50 ranked unique genes using both the standard GO and GO Slim biological process sets from PantherDB.org. Detailed specifications of the analysis can be found in **Supp. Table 2**. Top of hierarchy terms were used. Terms describing similar process such as ion transport-related were removed for clarity.

### co-IP and western blots of potential interactors

HEK293T cells were grown on 15 cm treated dishes in 20 ml DMEM + 10% FBS. Separate cultures of cells (not co-transfected) were transfected with either 3xFLAG- or myc-tagged proteins (pCMV vector). Cells were washed and resuspended in 5 ml cold PBS supplemented with 1 mM phenylmethanesulfonyl fluoride, 583 µM pepstatin A, 762 µM leupeptin, 10.6 mM Nα-Tosyl-L-arginine methyl ester HCl, 10.8 mM tosyl-lysine-chloromethylketone HCl, 11.3 mM Nα-benzoyl-L-arginine methyl ester carbonate, 200 µM soybean trypsin inhibitor, pelleted, and frozen to lyse. Thaw and resuspend FLAG-tagged expressing cells in 1 ml and myc-prey cells in 0.5 ml cold PBS with protease inhibitors. Pass 10 times through 27-gauge syringe. For membrane proteins also add 0.3% Triton-X and incubate at 4°C for 10 mins while rotating. Centrifuge 10 min at 4°C 16,000 x g. Incubate FLAG-tag supernatant with 120 µl of 25% slurry of anti-DYKDDDDK (anti-FLAG) magnetic agarose beads (Pierce A36797), and incubate at 4°C for 1 hour. Wash 3X with 1 ml cold supplemented PBS using magnetic rack. Retain aliquots of myc-tagged input proteins for western blot analysis (15 µl + 15 µl H_2_O + 10 µl 5X gel loading dye). Combine myc-tag protein supernatent with FLAG-bound magnetic beads. Resuspend and take aliquot for FLAG-tagged input protein for western blot (5 µl + 15 µl H_2_O + 5 µl 5X gel loading dye). Add 100 µM NH_3_^+^-SAGRRFIVSPVPE-COO^-^ blocking peptide diluted in 25 mM Tris-HCl pH 7.75, 125 mM NaCl to one of two samples per prey protein. Incubate at 4°C for 1 hour. Great care must be taken to perform the 3X washes as fast as possible due to the transient nature of some of the interactions (<30 sec for all three washes). Remove excess liquid from the side of the tube and on cap by aspiration. Due to large amount of FLAG relative to myc proteins, which may interfere with blots, myc proteins are eluted from the beads with 40 µl 500 µM blocking peptide. Elute with 40 µl 500 mM blocking peptide in PBS by gently vortexing and incubate on ice for 30 mins. Centrifuge and carefully collect 40 ul of supernatant without disturbing beads and add 15 µl 5X gel loading dye to obtain myc protein co-IP sample. To obtain FLAG protein IP samples add 50 µl 1:1 water:5X gel loading dye to beads and vortex thoroughly. Heat all gel samples at 65°C for 10 mins. Dilute FLAG-SPAK bait protein IP samples an additional 10X in 1X gel loading dye. Load bait and prey proteins on separate gels (Bio-Rad 4-20% precast gels, 10 well, 50 µl well volume cat. #4568094). FLAG-SPAK input: 5 µl, FLAG-SPAK IP: 5 µl, myc-prey input: 20 µl, myc-prey co-IP: 27 µl. Transfer all samples to nitrocellulose membrane 30 min, 750 mA at 4°C. Membranes incubated with 1:1 LI-COR blocking buffer:TBST (TBS + 0.1% Tween) overnight at 4°C. FLAG blots were incubated with 1:2500 Anti-FLAG M2 antibody (mouse, Sigma #F1804) in 1:1 blocking buffer:TBST. Myc blots were incubated with myc antibody from clone 9E10 (mouse, specific antibody no longer commercially available). Blots were washed 3 x 20 ml with TBST then incubated with 1:5000 IRDye 680 goat anti-Mouse secondary antibody (LI-COR # 926-68070). Blots were washed 3 x 20 ml with TBST then imaged using LI-COR imaging system.

### Kinase Assay

15 µM mouse His_6_-SPAK 62-556 T243E, 15 µM Mo25α, 5 µM MBP-Cavin1-His_6_ (or MBP tag alone, maltose binding protein, and 100 µM CCT blocking peptide were pre-incubated at room temperature for 15 minutes in 20 mM HEPES pH 7.5 150 mM NaCl (or supplemented with NaCl to bring to 300 mM or 500 mM), 1 mM DTT prior to addition of 50 µM ATP, 15 mM MgCl_2_, and [γ-32P]ATP. Reaction occurred at room temperature for 10 mins prior to addition of 5X Laemmli sample buffer and heating at 100 °C for 5 minutes to stop reaction. Samples were run on a 4-20% SDS-PAGE gel.

### HDMEC culture

Human dermal microvascular endothelial cells (HDMEC, PCS-110-010) obtained from ATCC (Manassas, VA, USA) were maintained in MCDB 131 medium (15100CV, Corning, Manassas, VA, USA) with 10 % FBS (F0926, Sigma-Aldrich),10mM L-glutamine (25005CI, Corning), 10 ng/ml epidermal growth factor (PHG0311, Thermo Fisher Scientific), 1 ug/mL hydrocortisone (H0888, Sigma-Aldrich), and 1 % penicillin/streptomycin (SV30010, Thermo Fisher Scientific). All cells were cultured at 37 °C with 95% humidity and 5% CO_2_.

### Antibodies

Unless indicated elsewhere in methods antibodies were obtained from the following sources: WNK1, Cobb lab Q2561 (Xu et al., 2000); Cavin1 #69036, OSR1 #3729, and GAPDH #97166L from Cell Signaling Technology (Beverly, MA, USA); phospho-SPAK Ser373 / phospho-OSR1 Ser325 07-2273 from Millipore Sigma (Darmstadt, Germany); phospho-Cavin1 Tyr156 PA5-99575 from Thermo Fisher Scientific (Waltham, MA, USA); Actin sc-8432 from Santa Cruz Biotechnology (Santa Cruz, CA, USA); TSC22D1 A303-582A from Bethyl Labs (Montgomery, TX, USA).

### siRNA knockdown

20 nM oligonucleotides encoding siRNA for control (Thermo Fisher Scientific, 4390844), TSC22D1 (Thermo Fisher Scientific #4392420 ID #S16891), and WNK1 (5’ CAGACAGUGCAGUAUUCACTT 3’) were transfected into HDMEC using Lipofectamine RNAiMax reagent (Thermo Fisher Scientific, 13778150). After 48 (TSC22D1) or 72 (WNK1) hours of transfection, cells were provided with their respective treatments and were then harvested in 1X SDS buffer (0.05% bromophenol blue, 0.1M DTT, 10% glycerol, 2% SDS, and 0.05M Tris-Cl) with 5% β-mercaptoethanol.

### Reverse transcription and qPCR

RNA from HDMEC was extracted using PureLink RNA kit (Thermo Fisher Scientific, 12183018A). RNA concentration was measured and 1 μg was used to synthesize cDNA using iScript™ Reverse Transcription Supermix (Bio-Rad, 1708891) as per manufacturer’s instructions. The resultant cDNA was diluted 1:10 in nuclease free water and the concentration was measured. The qPCR reactions with the cDNA and appropriate forward and reverse primers were set up using iTaq Universal SYBR® Green Supermix (Bio-Rad, 1725121) as per manufacturer’s instructions. The qPCR cycle is as follows: 95°C for 5 minutes, then 95°C for 10 seconds and 55°C for 30 seconds for 40 cycles. Custom oligos were purchased from Thermo Fisher Scientific. TSC22D1-isoform 3 (Forward Primer: 5’-ACTGCAGCCTACTCCTTGCT-3’; Reverse Primer: 5’-GAGTTCTTCTCAAGCAGCTC-3’). TSC22D3 (Forward Primer: 5’-CATGGAGGTGGCGGTCTATC-3’; Reverse Primer: 5’-CACCTCCTCTCTCACAGCGT-3’). TSC22D3-L isoform (Forward Primer: 5’-ACCGCAACATAGACCAGACC-3’; Reverse Primer: 5’-CACAGCGTACATCAGGTGGT-3’). TSC22D2 (Forward Primer: GCAGCGTCCTGACTAGATCC; Reverse Primer: GAGCTGTCAGTTCCCGACTC). 18s (Forward Primer: GTAACCCGTTGAACCCCATT; Reverse Primer: CCATCCAATCGGTAGTAGCG).

### Stable cell line generation and immunofluorescence microscopy

Lentiviral vectors containing C-terminal mNeonGreen tag were purchased from VectorBuilder, and TSC22D1 (long isoform) was inserted to generate TSC22D1-mNeonGreen. A549 and HEK293T cells were transfected using Lipofectamine 3000 according to manufacturer’s instructions. The Lipofectamine 3000:P3000:DNA ratio was 2.5 µl:2 µl:1 µg. Cells expressing the plasmids were selected by growing them in media containing 1 or 2 µg/µl puromycin (A549 or HEK293T, respectively). Then cells were trypsinized and sorted based on the fluorescent protein expression by flow cytometry. Immunofluorescence microscopy and image analysis was performed as described in (Gallolu Kankanamalage et al., 2016) with no 3D deconvolution.

## Supporting information

Supplemental Table 1 - CCT Interaction Predictions

Supplemental Table 2 - GO Analysis

## Acknowledgments

We thank other members of the Cobb laboratory, the Albanesi (Department of Pharmacology) and Goldsmith (Biophysics) labs and Kate Luby-Phelps [Department of Cell Biology and Live Cell Imaging Facility, University of Texas Southwestern (UT Southwestern)] for their suggestions about this project and Dionne Ware for administrative support. We thank the lab of Liang Tong (Columbia University) for working with us on one of the predicted hits. We would also like to thank three students who were involved in early stages of this work: Ashesh T.Triveda who was in the UTSW Summer Undergraduate Research Fellowship Program, Sakina Plumber who was in the UTSW-UT Dallas Green Fellows Program, and Jonathan Zijiang Yang who was in the UTSW STARS Summer Research Program. This project was supported by Welch Foundation Grant I1243 (to M.H.C.), National Institutes of Health Grants R01HL147661 (to M.H.C.), and National Cancer Institute Grant P30CA142543 to the Harold C. Simmons Comprehensive Cancer Center for partial support of the Live Cell Imaging Facility. J.J. was supported in part by the CPRIT training grant RP210041 and A.B.J. was supported in part by NIH K99AG075161-01A1.

## Notes

### Competing Interest Statement

The authors have declared no competing interest.

