## Supplemental Figures for "Predictive and Experimental Motif Interaction Analysis Identifies Functions of the WNK-OSR1/SPAK Pathway"

**TITLE**

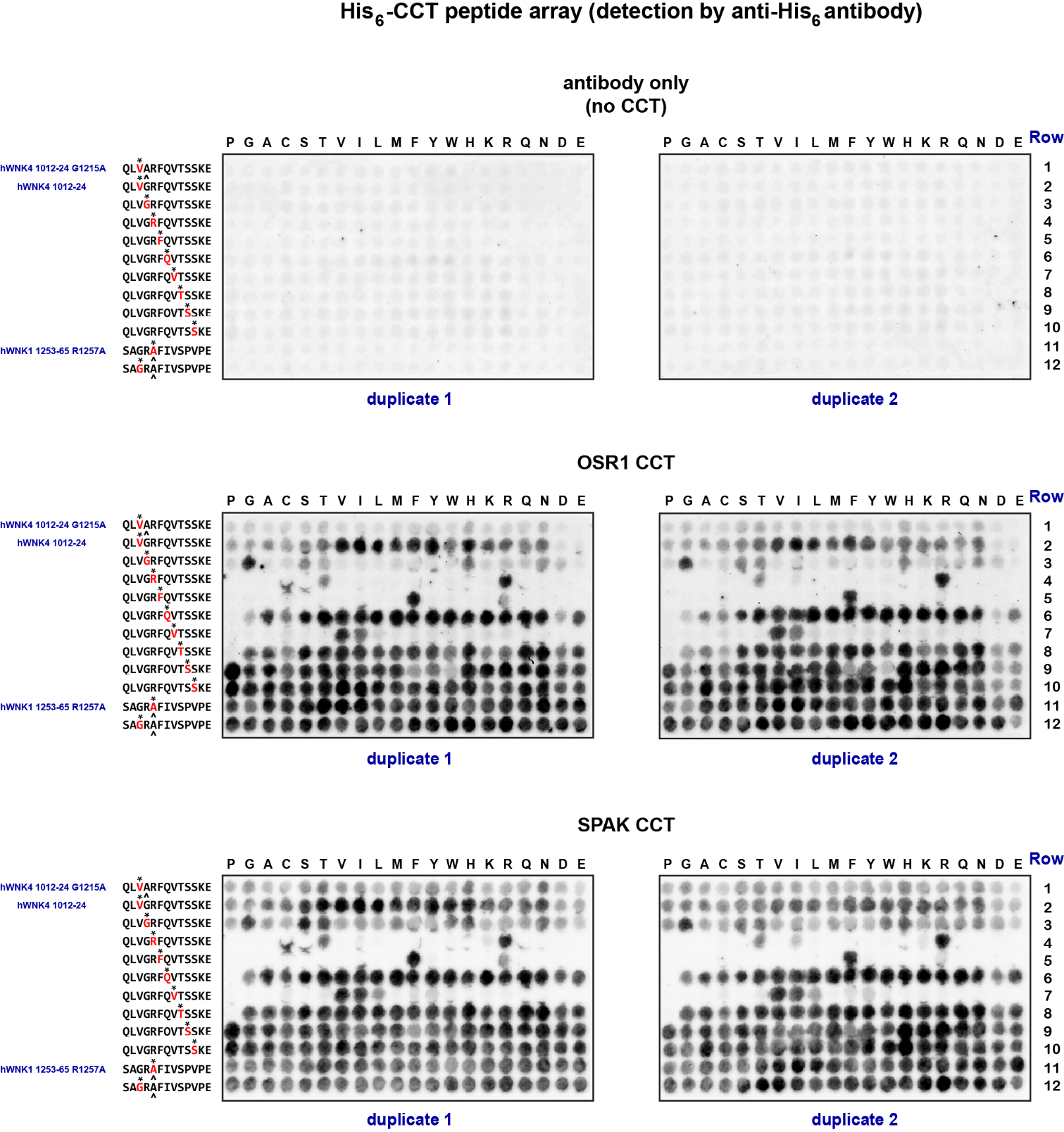

**Supp. Figure 1. Peptide array dot blots.** Experimental setup diagram can be found in Figure 1C. His_6_-SPAK and His_6_-OSR1 CCT domains were incubated with dot blot membranes containing indicated peptides. Membranes were washed and probed using anti-His_6_ primary and fluorescently labeled secondary antibodies. Imaging was done on LI-COR. Quantification of dot intensities was done using LI-COR Odyssey 3.0 software. Red letters with an asterisk (*) on left of blots indicates amino acid mutated in each row. Caret (^) under amino acid indicates amino acid substitution in base peptide that differs from the wild type sequence. Letters on top of blots indicate what the amino acid was mutated to. Antibody only blot was used as a control to detect unwanted interactions between peptides and antibodies.

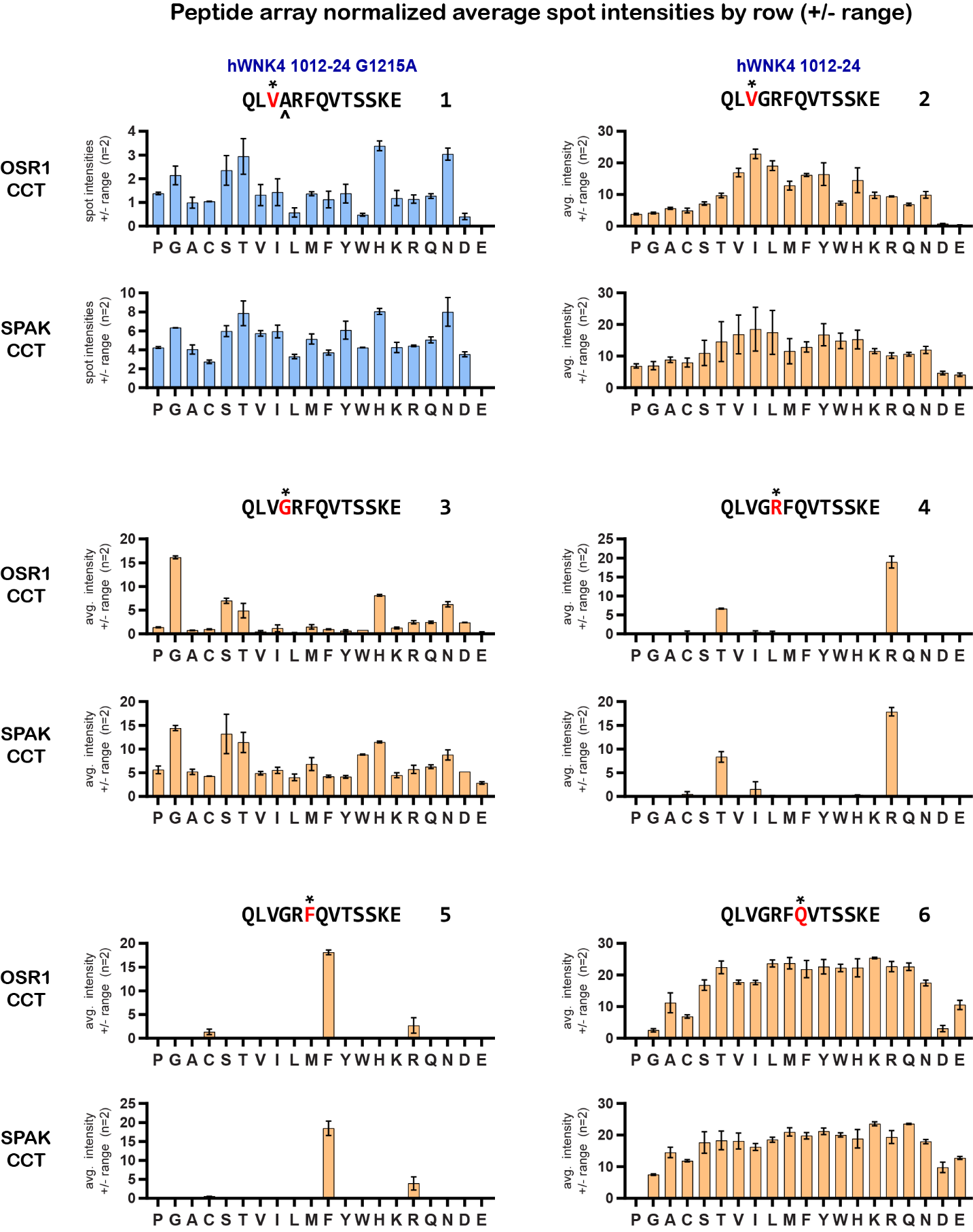

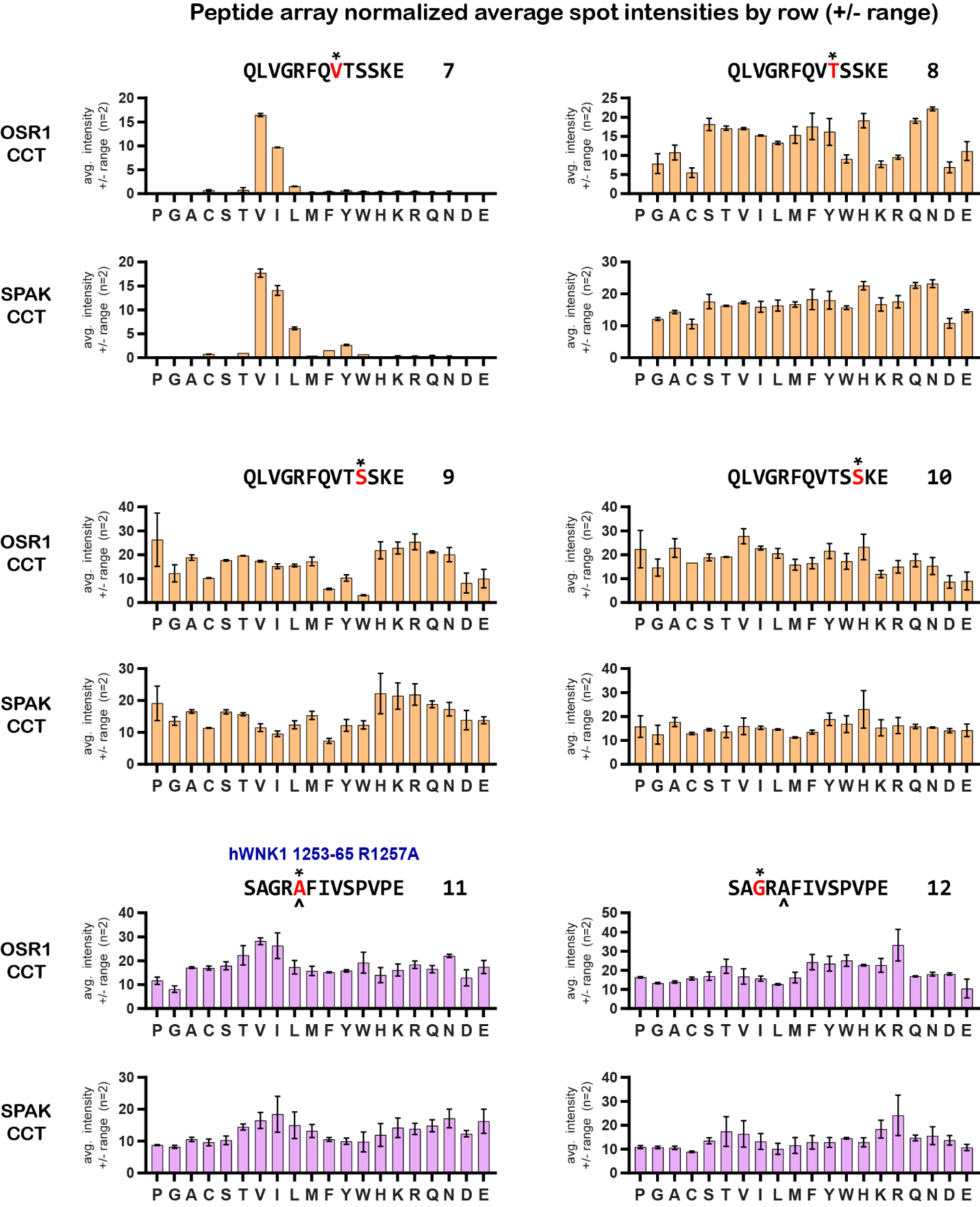

**Supp. Figure 2. Quantification of peptide array blots.** Average spot intensities for blots found in Supp. Fig. 1. Blots were normalized by average spot intensity across the entire blot and values for antibody only blot were subtracted. Bars represent range (n=2). Red letters with an asterisk (*) indicate amino acids mutated. Caret (^) under amino acid indicates amino acid substitution in base peptide that differs from the wild type sequence. Letters on x-axis indicate what the amino acid was mutated to.

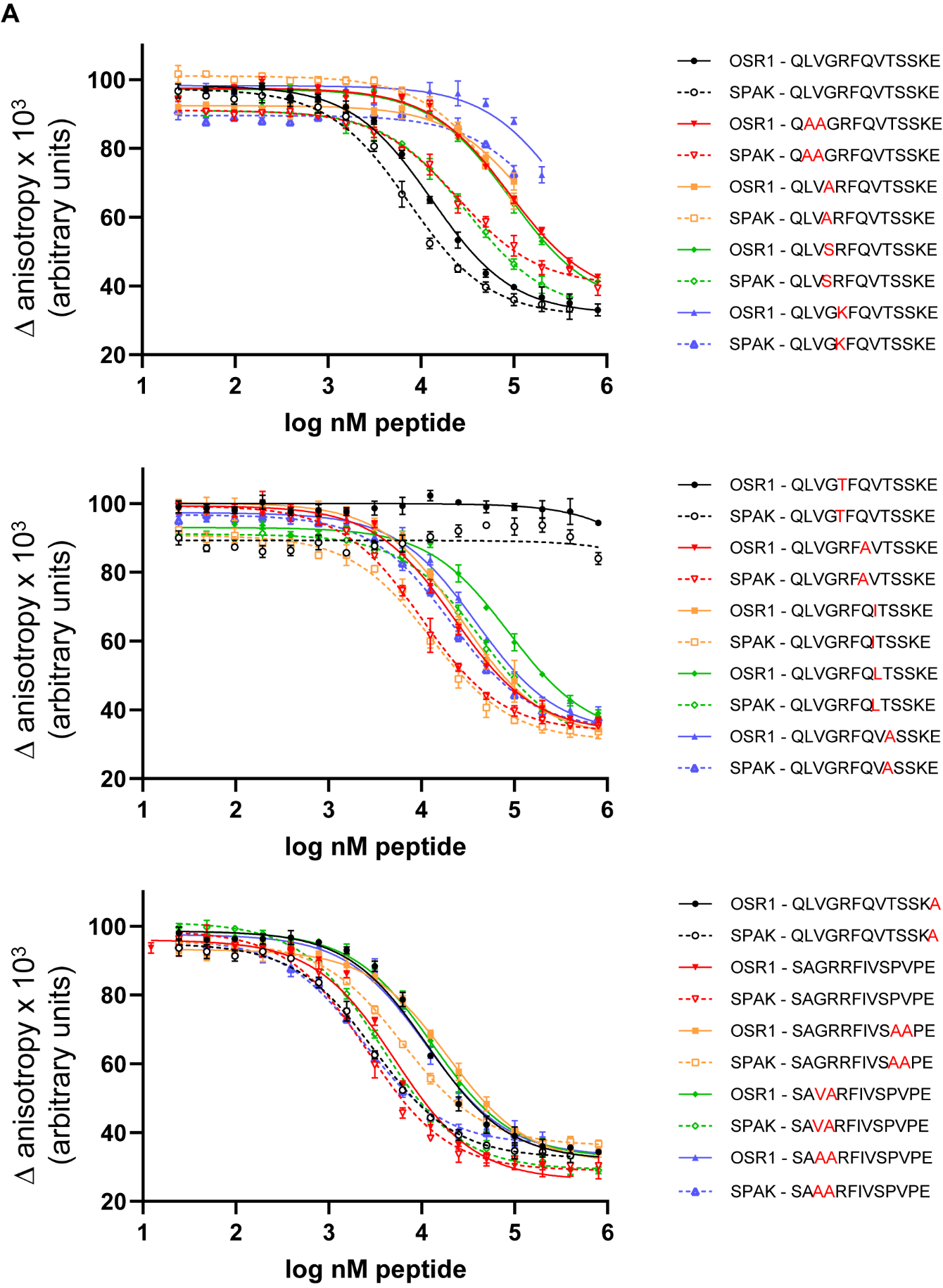

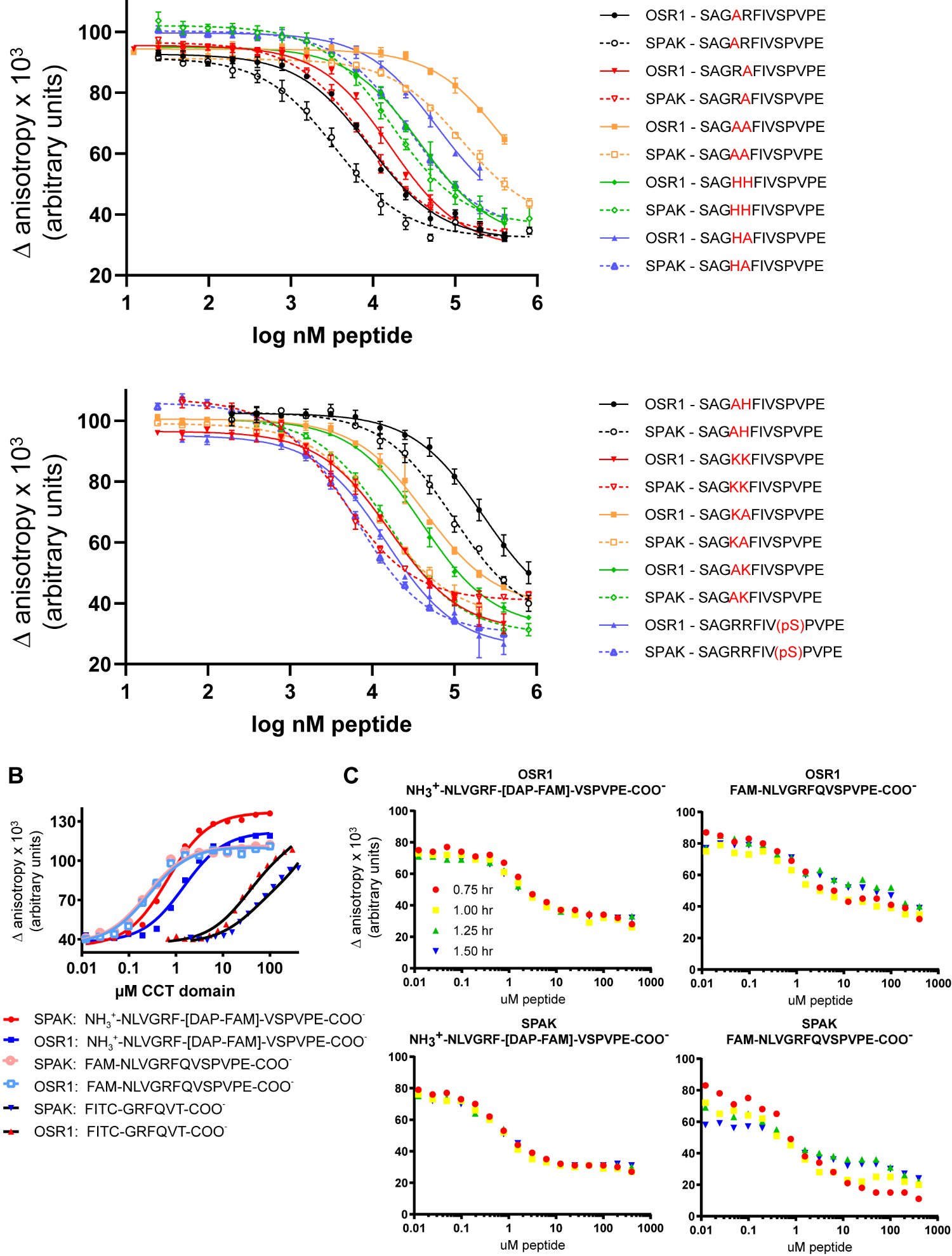

**Supp. Figure 3. Fitted curves for fluorescence anisotropy peptide competition. (A)** Unlabeled peptides displace labeled peptide (NH_3_^+^-NLVGRF-[DAP-FAM]-VSPVPE-COO^−^) [diaminopropionic acid (DAP)]. Labeled peptide held constant at 25 nM, SPAK CCT and OSR1 CCT are constant at 1.5 μM and 3.0 μM, respectively (n=3). Red letters indicate site of mutation. **(B)** Analysis of different labeled probe peptides for use in peptide FP competition assays. Varying concentrations of SPAK and OSR1 CCT domains were titrated. K_d_ values were: SPAK: NH_3_­^+^-NLVGRF-[DAP-FAM]-VSPVPE-COO^-^, 0.67 µM; OSR1: NH_3_­^+^-NLVGRF-[DAP-FAM]-VSPVPE-COO^-^, 1.5 µM; SPAK: FAM-NLVGRFQVSPVPE-COO^-^, 0.24 µM; OSR1: FAM-NLVGRFQVSPVPE-COO^-^, 0.26 µM; SPAK: FITC-GRFQVT-COO^-^, 34 µM; OSR1: FITC-GRFQVT-COO^-^, 49 µM. All probes held constant at 25 nM (n=1). **(C)** Stability of FP signal over time for indicated CCT domain and probe using QLVGRFQVTSSKA as the competing peptide. The higher affinity probe was not stable (n=1).

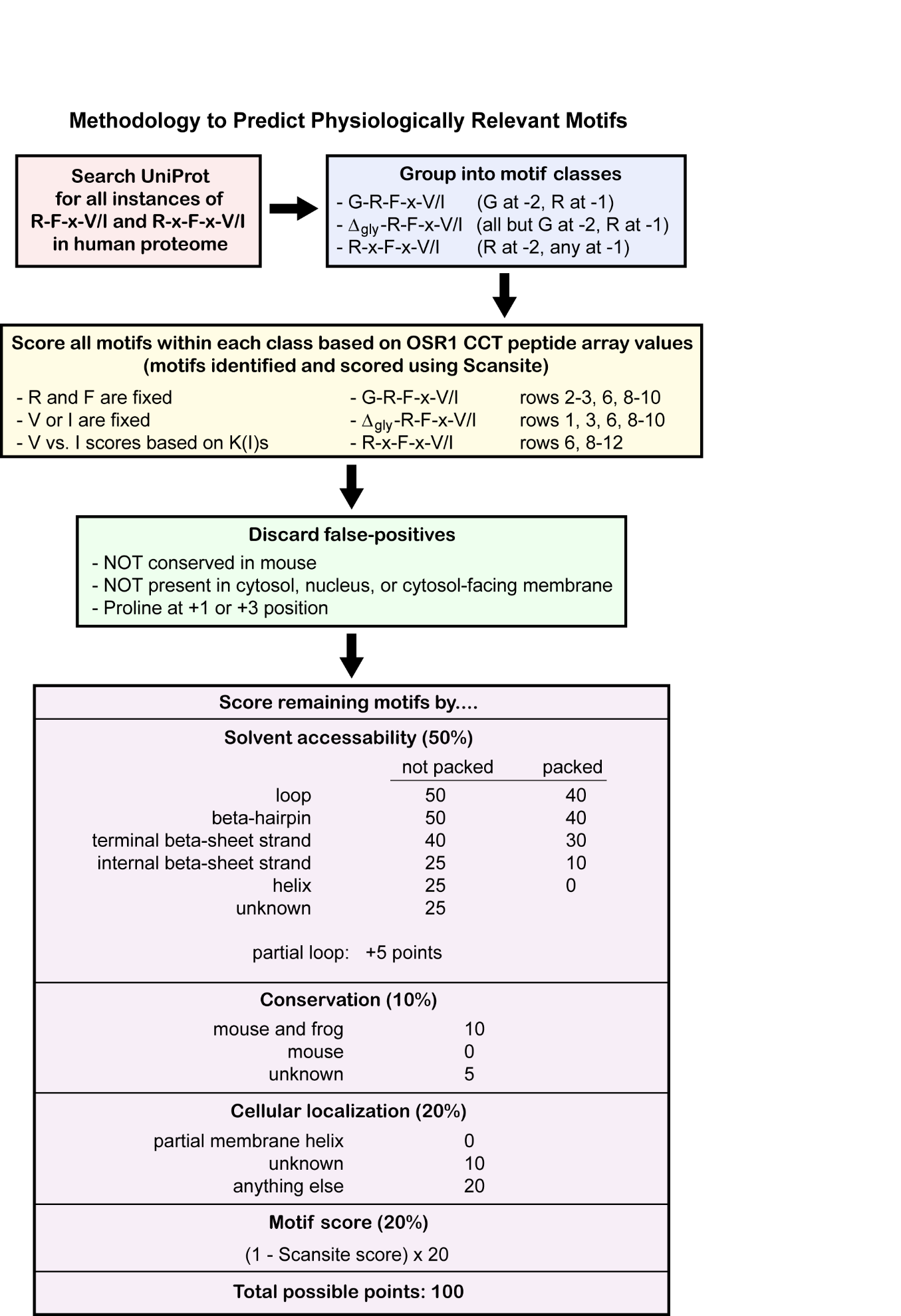

**Supp. Figure 4. Detailed flowchart for prediction analysis.** Elaboration of summary provided in Figure 2.

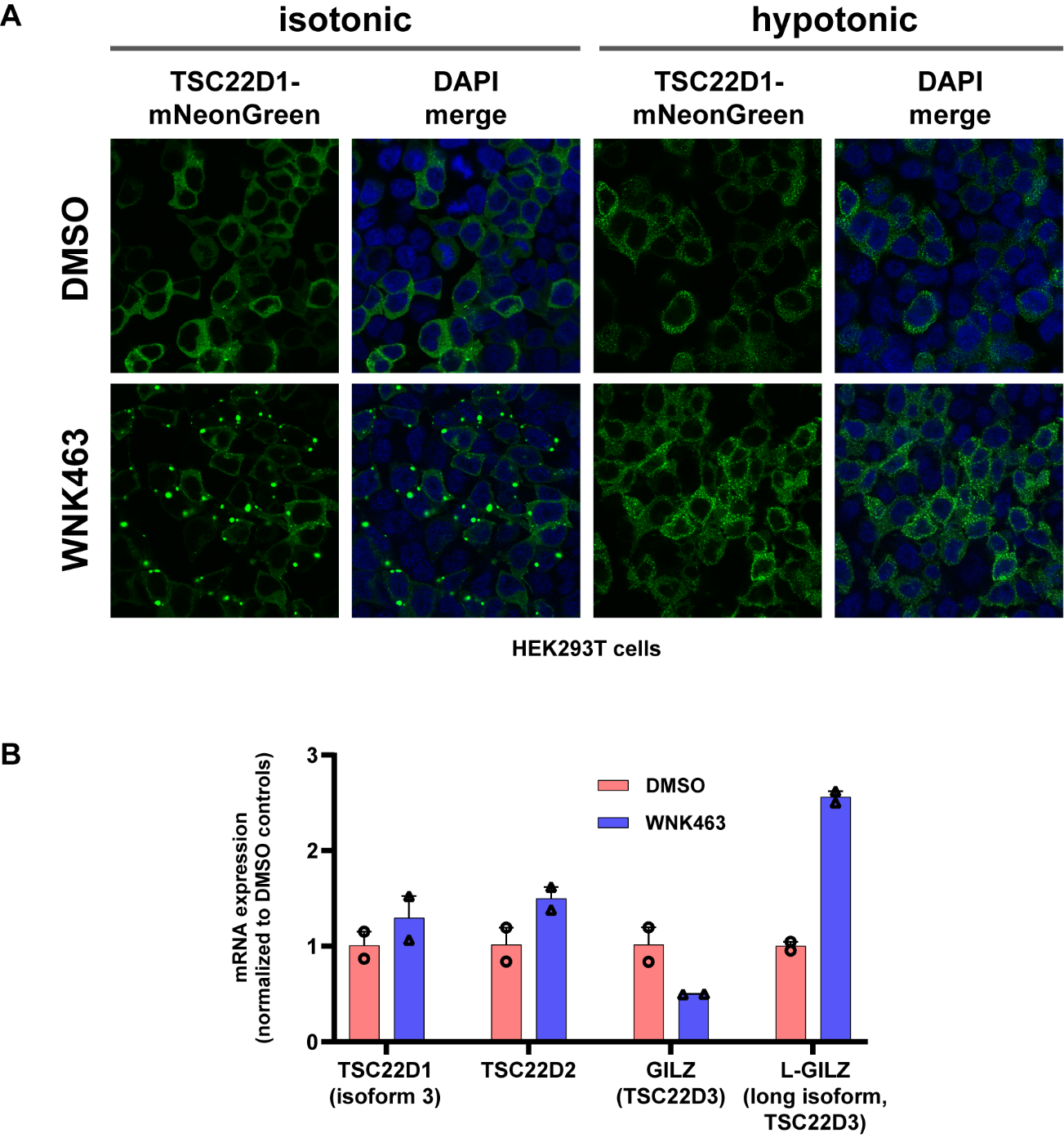

**Supp. Figure 5. WNK inhibitor affects TSC22D1 localization and expression. (A)** HEK293T cells stably expressing TSC22D1-mNeonGreen were treated with DMSO or 1 µM WNK463 for 24 hours. Additionally, hypotonic treatment was 1:1 DMEM media diluted with H_2_O for four hours (at end of 24 hour treatment with inhibitor). n=1. **(B)** Human dermal microvascular endothelial cells (HDMEC) were treated with DMSO or 1 µM WNK463 for 14 hours. Data represented as average with range from two replicates.

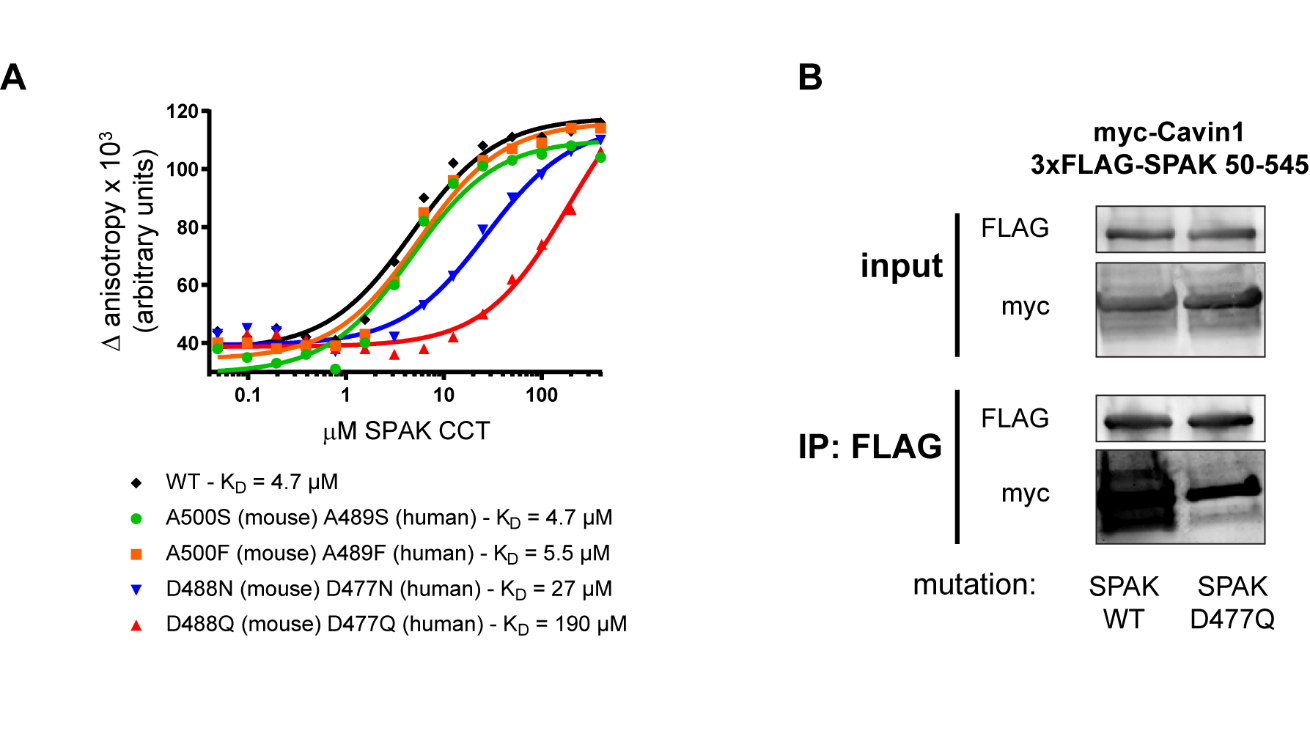

**Supp. Figure 6. Identification of CCT binding deficient mutants. (A)** Fluorescence anisotropy of labeled peptide (NH_3_^+^-NLVGRF-[DAP-FAM]-VSPVPE-COO^−^) [diaminopropionic acid (DAP)] in the presence of varying concentration of SPAK CCT domain. Probe kept constant at 25 nM. Mutation of CCT domain with both human and mouse numbering indicated along with measured K_D_ (n=1). **(B)** 3xFLAG-tagged SPAK 50-545 ± D477Q (binding-deficient mutant) was used to co-IP myc-Cavin1 in HEK293T cells (n=1).
